# A Cytoplasmic Index for Quantifying Immune-Related A-to-I RNA Editing

**DOI:** 10.64898/2025.12.03.692070

**Authors:** Roni Cohen-Fultheim, Itamar Twersky, Haim Krupkin, Shalom Hillel Roth, Erez Y. Levanon, Eli Eisenberg

**Affiliations:** Institute of Nanotechnology and Advanced Materials, Bar-Ilan University, Ramat Gan, 52900, Israel; Mina and Everard Goodman Faculty of Life Sciences, Bar-Ilan University, Ramat Gan, 52900, Israel; Raymond and Beverly Sackler School of Physics and Astronomy, Tel Aviv University, Tel Aviv, 6997801, Israel; Department of Genetics, Stanford University, Stanford, CA 94305, USA

**Author notes:** These authors contributed equally to this work (Roni Cohen-Fultheim, Itamar Twersky). These authors contributed equally to this work.

## Abstract

Distinguishing self from non-self is a major challenge for the immune system. Endogenous cytoplasmic double-stranded RNA (dsRNA) can mimic viral RNA and activate immune sensors like MDA5. ADAR1-mediated A-to-I editing disrupts base-pairing to suppress immunogenicity of these endogenous structures. Global editing indices are widely used to probe this crucial ADAR1 function. However, they are dominated by nuclear pre-mRNA edits with limited immune relevance. Here we present the Cytoplasmic Editing Index (CEI) that quantifies editing specifically within dsRNA structures in mature cytoplasmic transcripts, which carry higher immunological risk. Analyzing over 25,000 RNA-seq samples, we demonstrate CEI captures ADARp150 activity and outperforms the global editing index in terms of sensitivity and signal-to-noise, enabling sharper tissue-specific profiling, enhanced detection power of infection-induced editing changes, and stronger association with cancer prognoses. An open-source, cloud-native pipeline delivers end-to-end, reproducible analysis at very low cost, supporting immediate, scalable adoption.

**Micro-abstract:** The Cytoplasmic Editing Index (CEI) quantifies immune-relevant A-to-I RNA editing in inverted *Alu* clusters within 3′UTRs, capturing interferon-inducible ADAR1p150-dependent events. Analysis of >25,000 RNA-seq samples demonstrates CEI outperforms the global editing index in sensitivity and specificity, resolving tissue- and infection-linked editing patterns. An open-source, cloud-native pipeline enables scalable, low-cost deployment.

**Graphical Abstract:** 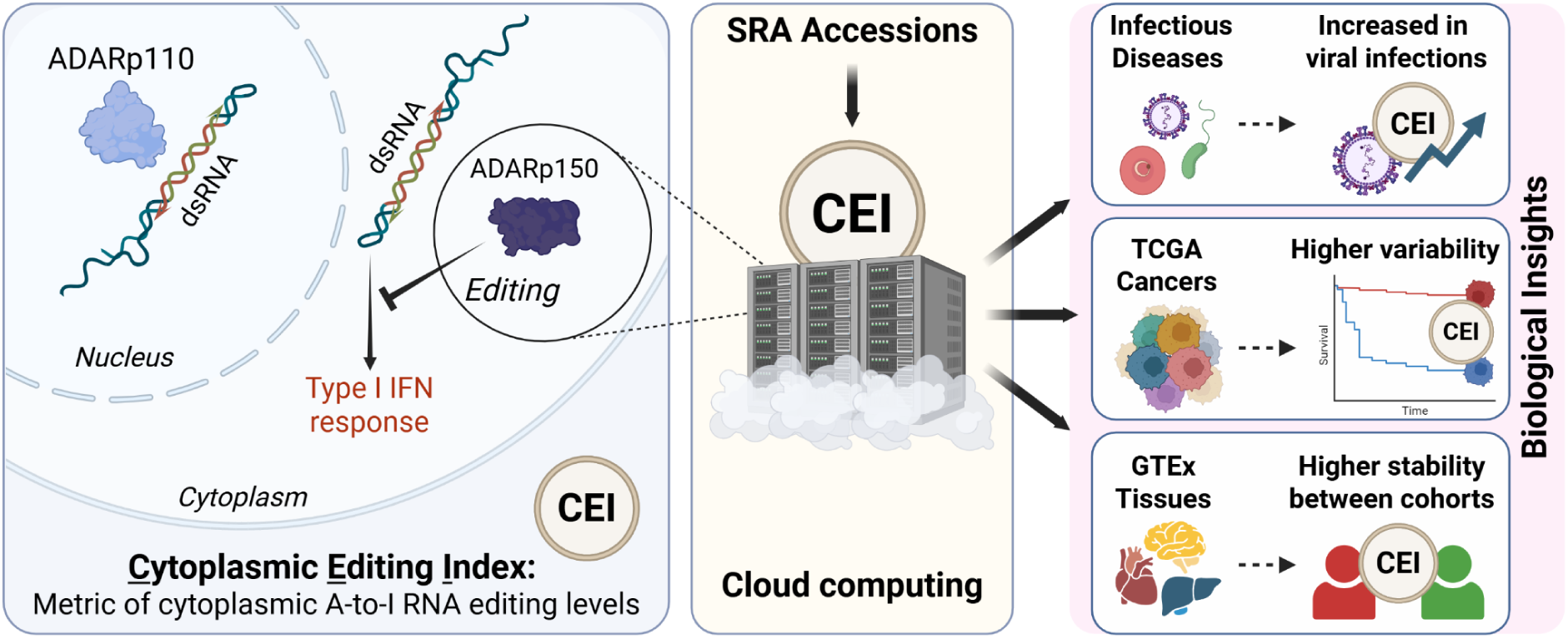

## Introduction

Adenosine-to-inosine (A-to-I) RNA editing is a widespread post-transcriptional modification carried out by adenosine deaminase acting on RNA (ADAR) family of enzymes that convert adenosines to inosines within double-stranded RNA (dsRNA)^1–3^. Among the ADAR enzymes, ADAR1 (ADAR) plays a central role in innate immune regulation. Endogenous dsRNAs that resemble viral RNA may be recognized by cytoplasmic RNA sensors and aberrantly activate antiviral pathways, most notably through the MDA5–MAVS axis and PKR, leading to type I interferon response^4–6^. Editing by ADAR1 disrupts the base-pairing of the immunogenic self-dsRNAs to the extent that they are no longer recognized by MDA5, or mark them otherwise, preventing inappropriate activation of the antiviral cellular immune system^7–10^. Aberrant editing has been associated with several autoimmune disorders^11–22^, and complete loss of ADAR1 is embryonic-lethal due to MDA5-mediated auto-immunity^7–9,23–25^.

In humans, the vast majority of editing events occur within primate-specific *Alu* retrotransposons^26–32^. These elements are abundant in primate genomes^33,34^, and frequently form long intramolecular dsRNA structures by pairing of neighboring inverted repeats^29,35^. Most *Alu* elements lie in intronic and intergenic regions^34,36^. Of the small fraction incorporated into mature mRNAs, *Alu* elements are especially enriched in noncoding regions, mainly the 3′ untranslated regions (3′UTRs). These elements may result in dsRNA structures that engage the cytoplasmic innate immune sensors such as MDA5^7–9,35^.

Given the essential function of A-to-I editing in preventing inappropriate immune activation, there is growing interest in accurate quantification of editing activity^32,37–44^. Current quantitative methods remain limited in interpretability and precision. The simplest method — counting edited sites — fails to account for the uneven distribution of *Alu* element coverage and the typically low editing frequency at individual sites within them. Accordingly, it is highly sensitive to sequencing depth and does not provide a robust estimate for enzymatic activity^45^. More advanced metrics, such as the *Alu* Editing Index (AEI)^39^, calculate a global score by averaging *Alu* editing frequencies across the human genome, weighted by expression levels. While these are useful for gauging overall ADAR activity^14,16,17,20,21,46–64^, they conflate signals from heterogeneous transcripts and subcellular compartments including intronic or nuclear-retained RNAs. The pre-mRNA structures are rarely seen in the cytoplasm^65^ and are therefore invisible to the cytoplasmic innate immune sensors. Previous analysis^66^ has found that only ∼0.15% of putative dsRNA structures in pre-mRNA exist in mature mRNAs. Furthermore, the vast majority of the global editing signal is due to a very large number of weakly edited *Alu* repeats^39,67^, each of which is not dramatically affected by editing, and unlikely to be immunogenic. Consequently, aggregate genome-wide editing indices are mostly dominated by weakly-edited nuclear structures, lack specificity for immune-relevant substrates and may be insensitive to the subtle dynamics of editing that regulate innate immunity.

To address this gap, we introduce a Cytoplasmic Editing Index (CEI): a biologically informed metric that specifically quantifies A-to-I editing events within putative cytoplasmic dsRNAs – clusters of inverted *Alu* element residing in 3′UTRs, which are expected to form long intramolecular dsRNA structures capable of activating cytoplasmic immune sensors^68^. Integrating both structural and spatial constraints, this metric provides focused measurement of editing in the most immunologically relevant substrates.

We demonstrate that the CEI provides enhanced sensitivity for detecting immunologically relevant editing changes and serves as a valuable tool for investigating RNA editing across diverse biological contexts. Benchmarking across over 25,000 RNA-seq samples from infectious disease and normal tissue datasets, we show that the CEI robustly captures editing dynamics under diverse conditions. To support broad adoption, we provide an accessible, cost-efficient cloud-computing framework enabling scalable and systematic quantification of immunogenic dsRNA editing across diverse experimental settings.

## Results

### A targeted and scalable index for cytoplasmic A-to-I RNA editing

To develop a cytoplasmic editing index that captures immunologically relevant dsRNA substrates, we first established quantitative criteria for identifying *Alu* elements with immunogenic potential (Fig. 1a). We defined inverted 3′UTR clusters as regions containing ≥2 *Alu* elements (each >200 bp) with at least one element oriented in each direction, enabling intramolecular dsRNA formation (Fig. 1d). The selected regions are both retained in cytoplasmic transcripts and capable of forming stable secondary structures detectable by cellular dsRNA sensors.

**Figure 1:**
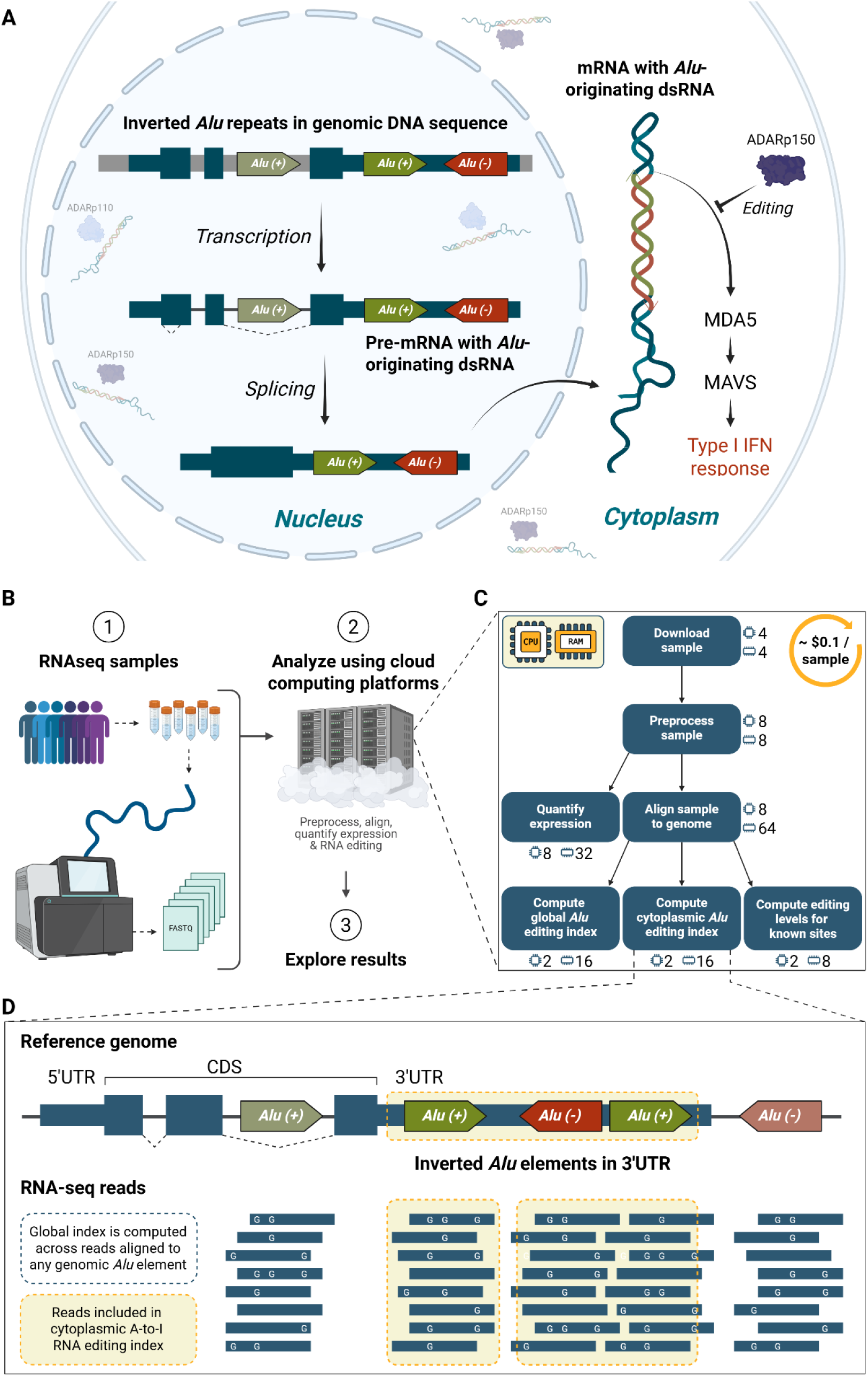
Cloud-based pipeline and biological rationale for quantifying cytoplasmic versus global A-to-I RNA editing. *a*, Biological rationale of the cytoplasmic editing index. For an RNA molecule to trigger immune activation, it must (1) localize to the cytoplasm and (2) adopt a double-stranded conformation therein. Following transcription and splicing, mRNAs are transported to the cytoplasm. Reversely oriented *Alu* repeats in the 3′ UTRs can fold into intramolecular double-stranded RNA (dsRNA) structures. When these cytoplasmic duplexes are insufficiently edited, they are recognized by cellular innate immune pathways and trigger type-I interferon responses. The cytoplasmic editing index specifically quantifies editing within these immunologically relevant dsRNA substrates. *b*, Cloud-based analysis workflow. RNA-seq samples are (1) retrieved, (2) processed through a modular cloud pipeline, and (3) analyzed for A-to-I editing patterns. The pipeline is designed for high-throughput processing and optimized for efficient deployment across diverse dataset sizes, with compatibility across multiple cloud-computing environments. *c*, Resource optimization and scalable processing. Each pipeline step operates with minimal computational requirements (virtual CPU and RAM allocations shown beside icons), with processing costs of approximately $0.10 per sample (for GCP and AWS platforms). The containerized Nextflow implementation enables portable deployment to any Nextflow-compatible cloud or local computing environment. Complete parameter specifications are detailed in Methods. *d*, Cytoplasmic vs. global editing index computation. The cytoplasmic editing index is calculated exclusively from RNA-seq reads mapped to inverted *Alu clusters* within 3′ UTRs (Methods). These clusters are likely to form long, intramolecular dsRNA structures that remain stable after splicing and nuclear export. By contrast, the global *Alu* editing index encompasses reads aligned to any genomic *Alu* element, regardless of orientation, size or genomic context.

Applying these criteria genome-wide (hg38), we identified 3,343 *Alu* elements across 895 genes within structured 3′UTR *Alu* clusters (see Methods). The Cytoplasmic Editing Index (CEI) is defined as the aggregate A-to-I editing level calculated exclusively across these immunologically relevant *Alu* elements. For comparison, we computed the global *Alu* editing index (AEI), which aggregates editing across 1,113,103 genomic *Alu* elements (merged contiguous elements of any length) irrespective of orientation or transcriptomic context^39^. The computational strategies underlying both indices are summarized in Fig. 1d.

To enable large-scale comparative analysis of cytoplasmic vs. global editing patterns, we developed a cost-efficient, modular, cloud-based computational pipeline that retrieves RNA-seq data, performs preprocessing, alignment, and quantifies expression and editing (Fig. 1b). The input file for this workflow is a list of SRA sample accession codes. Resource allocation is optimised for each step, resulting in a per-sample processing cost of ∼US $0.10 (Fig. 1c). We validated our approach by analyzing 36,334 publicly available RNA-seq samples from diverse contexts — infectious disease studies (15,649 samples), GTEx (9,602 samples) and TCGA (11,083 samples) — demonstrating robust scalability across heterogeneous datasets. All pipeline scripts and reference files are open-access to support reproducibility (Methods).

### Few *Alu* clusters contribute to the CEI

First, we characterize the 3,343 *Alu* elements within 3’UTRs contributing to the CEI, and their genomic architecture. We compared these to the 5,344 elements in tandem *Alu* clusters that contain one or more 3’UTR *Alu* elements all oriented in the same direction (Fig. 2a). The latter, including single-*Alu* clusters (Fig. 2d), are not expected to form cytoplasmic intramolecular dsRNAs. Furthermore, tandem clusters are present in ∼five-fold more genes than inverted clusters (Fig. 2f; 4,011 vs. 791 genes). Similarly, analysis of murine B1 and B2 repeat clusters in mouse 3′ UTRs show that tandem clusters dominate and inverted clusters are restricted to far fewer genes (Fig. 2e, g). These results suggest conserved evolutionary constraints against dsRNA-forming configurations in 3′ UTRs.

**Figure 2:**
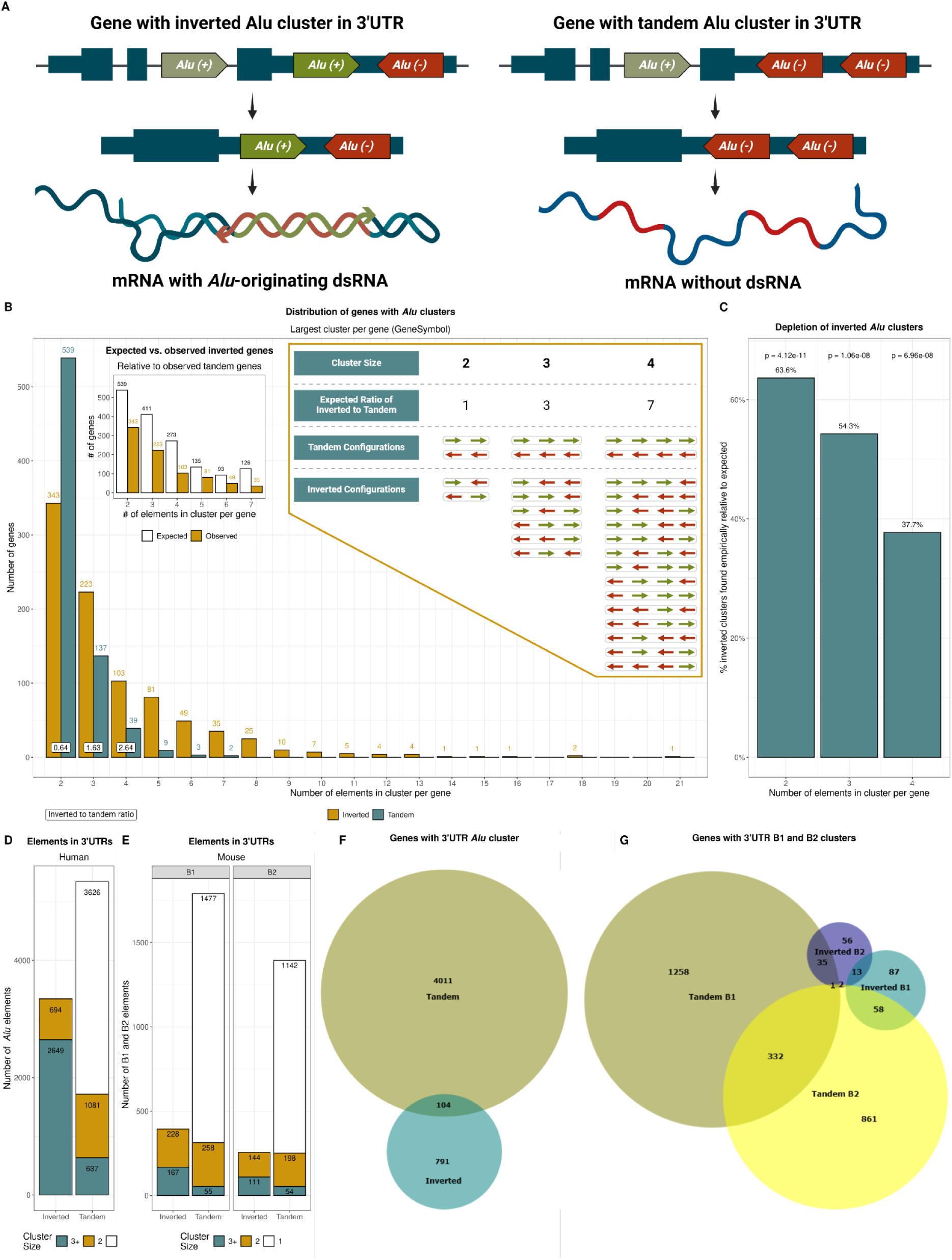
Structural constraint on inverted versus tandem *Alu* clusters in human 3′UTRs. *a*, Schematic of *Alu* configurations in 3′ UTRs. Inverted *Alu* clusters can fold into intramolecular dsRNA that can be edited by ADAR and can activate cytoplasmic RNA sensors if unedited, whereas tandem *Alu* clusters lack complementary elements and do not form dsRNA. *b*, Distribution of the largest *Alu* cluster per gene (cluster found in the transcript whose 3’UTR has the largest number of *Alu* elements among all gene’s transcripts), stratified by orientation. Right inset: possible tandem and inverted configurations for cluster sizes 2–4, and expected inverted:tandem ratios under a random-orientation model (see Methods). Left inset: observed vs. expected number of genes with inverted *Alu* clusters. Labels above each bar indicate gene counts per category; empirical inverted:tandem ratios (bottom, sizes 2–4) compare to right inset expectations. Inverted *Alu* clusters span a broader size range than tandem *Alu* clusters, reflecting the larger combinatorial space of mixed orientations. Nevertheless, inverted *Alu* clusters are consistently rarer than expected (panel c), indicating negative selection against dsRNA-forming configurations. *c*, Depletion of inverted clusters relative to the random model. Bars represent the observed proportion of inverted clusters divided by the proportion expected when orientations are assigned at equal probability (for *Alu* cluster sizes with > 10 genes per orientation; Methods; Supplementary Fig. S1). Values < 100% indicate underrepresentation, consistent with selective constraint against dsRNA-forming orientations. The proportion of observed inverted clusters declines with cluster size (χ² test, no multiple testing correction for the three tests; Supplementary Fig. S1). *d*, *e*, Orientation breakdown of *Alu* elements in human 3′ UTRs (*d*; 3,343 inverted and 5,344 tandem elements) and B1/B2 elements in mouse 3′ UTRs (*e*; 650 inverted and 3,184 tandem elements). Each element is assigned to the largest cluster size in which it participates within each orientation (singleton, pair or ≥3). Elements appearing in both orientation classes (for different transcript isoforms) are counted twice: 80 human *Alu* elements, 3 mouse B1 elements, and 1 mouse B2 element. In both species, tandem elements predominate and are overwhelmingly singletons, whereas inverted elements, though fewer overall, make the majority of multi-element clusters. *f*, Number of human genes harboring inverted and tandem 3′ UTR *Alu* clusters. There are 104 genes exhibiting both types, each in a different isoform. *g*, Same as *f* for mouse B1/B2 clusters (Euler diagram).

Examining tandem and inverted clusters abundance as a function of cluster size reveals depletion of inverted clusters compared to tandem clusters (Fig. 2b). For size-2 clusters, the number of tandem configurations exceeds that of inverted ones; for sizes ≥3, inverted clusters outnumber tandems (Fig. 2b), but not as much as expected in the absence of evolutionary purification (left inset, Fig. 2b). For a size-*N* cluster, there are 2^*N*^ orientation combinations, of which 2*^N^* − 2 are mixed-strand configurations associated with inverted clusters, compared to only two homogeneous-orientation ones associated with tandem clusters. Thus, a random, equally probable and uncorrelated assignment of orientations should result in an inverted:tandem ratio of 2^*N*−1^ − 1 for a cluster of size *N* (right inset, Fig. 2b). The observed ratio is well below this expectation (left inset, Fig. 2b), indicating strong depletion of dsRNA-forming configurations (Fig. 2c; Supplementary Fig. S1). This finding is consistent with previous reports on evolutionary selection against dsRNA-forming arrangements in mature transcripts^66^.

Similar results are obtained when the analysis is restricted to the longest 3′ UTR isoform, as well as for analyses of murine B1 and B2 repeat clusters in mouse 3′ UTRs (Supplementary Figs. S2; Supplementary Fig. S3).

We tested whether cluster architecture might be influenced by *Alu* evolutionary age or subfamily composition, but found only minor differences in sequence divergence and slight deviations from genome-wide proportions (Supplementary Fig. S1).

### Cytoplasmic index reflects ADARp150-dependent immune activation

ADAR1p150, the interferon-inducible cytoplasmic isoform of ADAR1^2,69^, plays a critical role in preventing cytoplasmic dsRNA from triggering innate immune responses. We thus asked whether the CEI, probing editing in *Alu* elements within inverted *Alu* clusters, exhibits elevated sensitivity to editing by ADAR1p150. We compared the CEI to two additional editing metrics: the conventional global *Alu* editing index (AEI), and a tandem editing index quantifying editing within tandem-oriented *Alu*s (negative control, see Methods). Analysis of knockout^10^ and reconstitution^70^ datasets revealed that the CEI was selectively sensitive to ADAR1p150 loss or reconstitution compared to the AEI, while the tandem index remained consistently low across all conditions.

As expected, ADAR1 knockout suppressed all editing indices; however, selective loss of ADAR1p150 reduced the CEI more strongly. Similarly, in the reconstitution system, ADAR1 knockdown reduced editing signals in both indices as well. Subsequent ADAR1p150 expression selectively restored the CEI almost to its wild-type levels, while global editing remained diminished (Fig. 3a).

**Figure 3:**
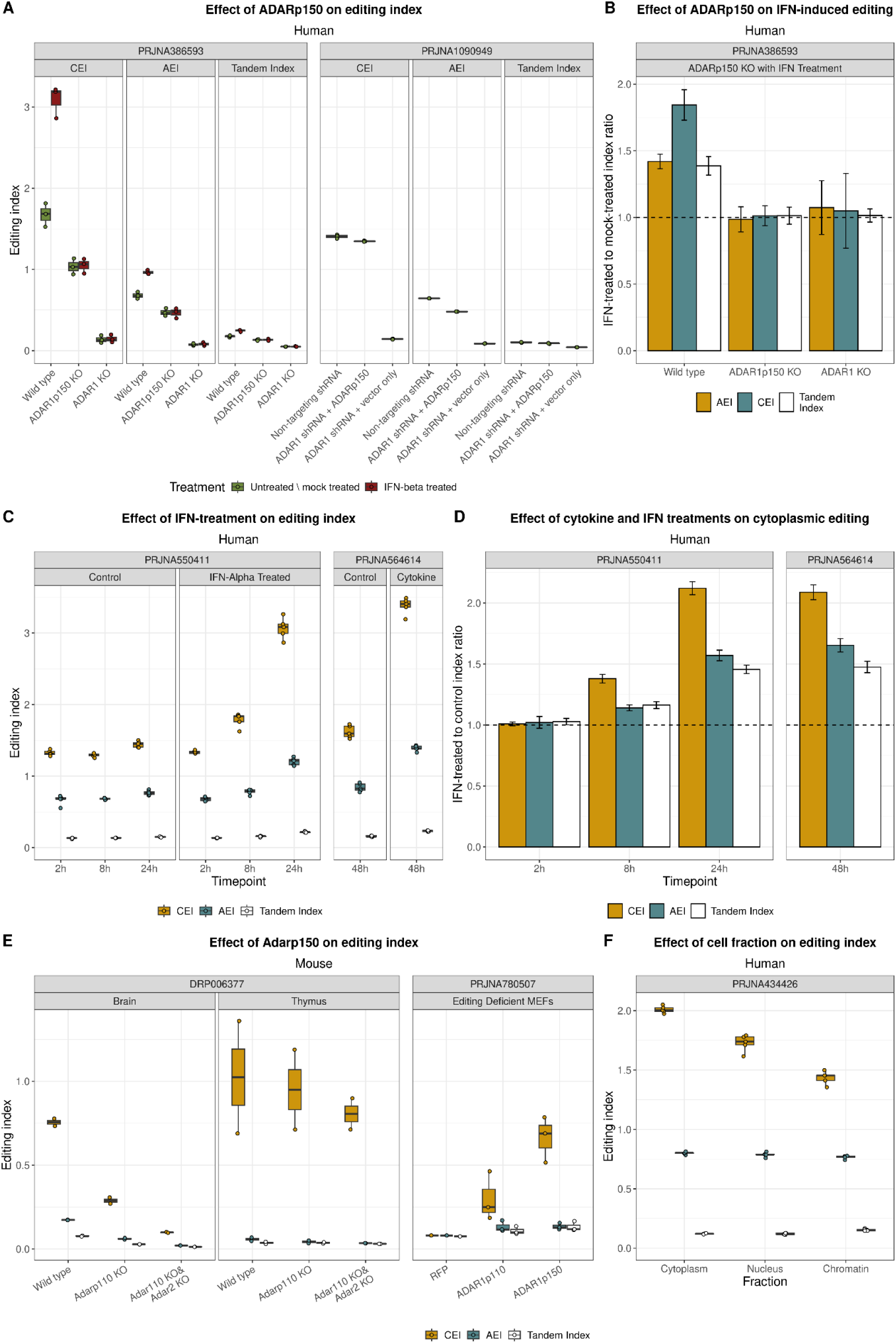
Biological perturbations of A-to-I RNA editing reveal differential sensitivity of CEI and AEI. *a*, Impact of ADAR1p150 loss and reconstitution on editing indices. Left^10^: HEK293 cells with ADAR1 knockdown followed by ADAR1p150 overexpression; right^70^: TBP-ARPE cells with ADAR1p150 knockout (KO) ± IFN-β treatment (each sample cultured and sequenced in triplicate). Loss of ADAR1 activity reduces global editing in both systems, while cytoplasmic editing is selectively impaired by ADAR1p150 absence and restored upon isoform presence, demonstrating that the cytoplasmic index specifically measures ADAR1p150-mediated activity. The tandem index (computed from exclusively tandem *Alu* elements, see Methods) remains near-zero across all genotypes, validating the assumption that these regions cannot form dsRNA. *b,* ADAR1p150 is required for IFN-β responsiveness. Bars show fold-change in mean editing (IFN-β treated versus untreated) from the ADAR1 KO experiment in panel *a* (mean ± s.d., n = 3). While IFN treatment enhances the cytoplasmic index in WT cells, this response is abolished in both ADAR1p150 KO and ADAR1 KO cells, confirming isoform specificity. Global and tandem indices show minimal IFN responsiveness in WT cells. *c,* Temporal dynamics of cytokine-induced editing in EndoC-◻H1 cells. Two independent time-course datasets (IFN-α, left^71^; mixed cytokines, right^72^) reveal index trajectories over 2–48 h. The cytoplasmic index rises earlier and more dramatically than the global index, while the tandem index remains consistently low and largely unaffected by cytokine treatment. *d,* Fold-induction for datasets shown in panel *c* (mean ± s.d., n = 5). The CEI consistently exhibits greater dynamic induction compared to the AEI. Apparent fold-changes in the tandem index do not reflect biologically meaningful induction, as the absolute editing levels remain near baseline (panel *c*). *e*, Species conservation of ADAR1p150 dependence. Left^74^, tissue-specific patterns: brain (predominantly ADAR1p110-expressing) shows marked reduction of CEI and AEI upon ADAR1p110 knockout, with a further decrease in the ADAR1p110/ADAR2 double knockout. In contrast, thymus (predominantly ADAR1p150-expressing) maintains high editing across genotypes. Right^75^, rescue experiments: editing-deficient MEFs (ADAR1⁻/⁻; ADARB1⁻/⁻; Gria2ᴿ/ᴿ) show no editing when expressing RFP controls, modest CEI restoration with ADAR1p110, and complete rescue with ADAR1p150. In both contexts, AEI and tandem indices remain consistently flat. *f,* Subcellular specificity. Analysis of subcellular fractions from HeLa cells^76^ reveals highest CEI in the cytoplasm, decreasing progressively in nuclear and chromatin fractions, whereas AEI and tandem indices remain relatively constant across compartments.

In addition, the knockout dataset included IFN-β stimulation, enabling assessment of interferon-induced editing responses across genotypes. In wild-type cells, IFN-β treatment induced a marked increase in the CEI compared to untreated controls, while the AEI showed no significant change and the tandem index remained largely unresponsive. IFN-induced enhancement of cytoplasmic editing was absent in both complete ADAR1 knockout and selective ADAR1p150 knockout cells (Fig. 3b), confirming that cytoplasmic editing induced by interferon specifically requires ADAR1p150.

Time-course analyses in two independent datasets^71,72^ revealed that CEI rose faster and higher than AEI following IFN-α or cytokine treatment, whereas the tandem index remained largely unaffected (Fig. 3c,d). Similarly, the cytoplasmic index exhibits preferential responsiveness to cytokines in human islets^71,73^ (Supplementary Fig. S4). The contrasting behavior of inverted versus tandem *Alu*s — despite both being part of cytoplasmic 3′ UTRs — demonstrates that structural architecture, not subcellular context alone, determines immune-responsive editing activity.

Analysis of mouse tissues^74^ and embryonic fibroblasts^75^ demonstrated evolutionary conservation of isoform-specific cytoplasmic editing. In mouse thymus, predominantly expressing ADAR1p150^74^, cytoplasmic editing does not change much upon knockout of ADAR1p110 and ADAR2. Conversely, in mouse brain tissue, primarily expressing ADAR1p110^74^, ADAR1p110 knockout causes marked CEI reduction; combined ADAR1p110/ADAR2 knockout further diminishes this effect. In editing-deficient fibroblasts (ADAR1⁻/⁻; ADARB1⁻/⁻; Gria2ᴿ/ᴿ), cytoplasmic editing was rescued robustly by ADAR1p150, partially by ADAR1p110, and not at all by an RFP control. Across these conditions, the effect on AEI and tandem index is limited, confirming the isoform specificity of cytoplasmic editing (Fig. 3e).

Direct assessment of subcellular localization further supported the cytoplasmic specificity of the CEI. Fractionation experiments in HeLa cells^76^ revealed highest CEI in cytoplasmic fractions, intermediate in nuclear fractions, and lowest in chromatin fractions, whereas the AEI and tandem indices presented negligible compartmental variation (Fig. 3f). Thus, CEI demonstrates isoform specificity absent in conventional editing indices, with particular utility for monitoring immune activation and infection responses

Collectively, these findings establish CEI as a ADAR1p150-sensitive, biologically grounded, and computationally efficient metric that specifically captures editing activity in immunologically relevant dsRNA substrates.

### Cytoplasmic editing index provides superior signal, specificity and computational performance

We next evaluated the technical performance of the CEI relative to the widely used global *Alu* editing index (AEI)^39^. Leveraging our standardized cloud-based computational pipeline (Fig. 1b,c), we quantified both indices across 15,649 publicly available RNA-seq samples from 55 infectious-disease datasets, enabling consistent large-scale benchmarking (Fig. 1b,c; Fig. 5a; Methods). We compared editing signal, signal-to-noise ratio, and mean coverage per editable site to assess the relative advantages of the two indices.

For both indices, A-to-G mismatches consistently ranked first in frequency (>99% of samples), with C-to-T second (>93%) (Supplementary Fig. S5). The values reported for the two indices were correlated across samples, but CEI consistently outperformed AEI across multiple metrics. CEI delivered a ∼2.6-fold higher editing signal than AEI (ordinary least squares slope = 2.57; Fig. 3a) and exhibits a 2.15-fold higher signal-to-noise ratio (Fig. 3b), reflecting improved sensitivity and specificity for A-to-I events. This enhanced performance stems partly from CEI’s ∼16-fold greater average coverage per editable site (slope = 16.5; Fig. 3c), reflecting its restriction to mature, cytoplasmic transcripts. These gains were obtained without increasing variability across datasets (Fig. 3d), underscoring CEI’s efficiency and robustness for large-scale RNA editing analysis. Given that ∼0.3% of the genomic *Alu* elements are included in the tissue-specific index, their coverage is ∼16.5-fold enriched, and their editing ∼2.15-fold enhanced, we estimate that approximately 11% of the A-to-G mismatches contributing to the AEI are captured in the CEI.

The distribution of the noise per sample is similar for both indices, with a slight advantage to the global index (Fig. 4e). In addition, the cytoplasmic index remained robust to sub-optimal mapping conditions (Supplementary Fig. S5). Finally, CEI required substantially lower compute time and memory usage than AEI (Fig. 4f,g), highlighting its efficiency. These advantages are recapitulated in 9,602 Genotype-Tissue Expression^77^ (GTEx) samples (Supplementary Fig. S6).

**Figure 4:**
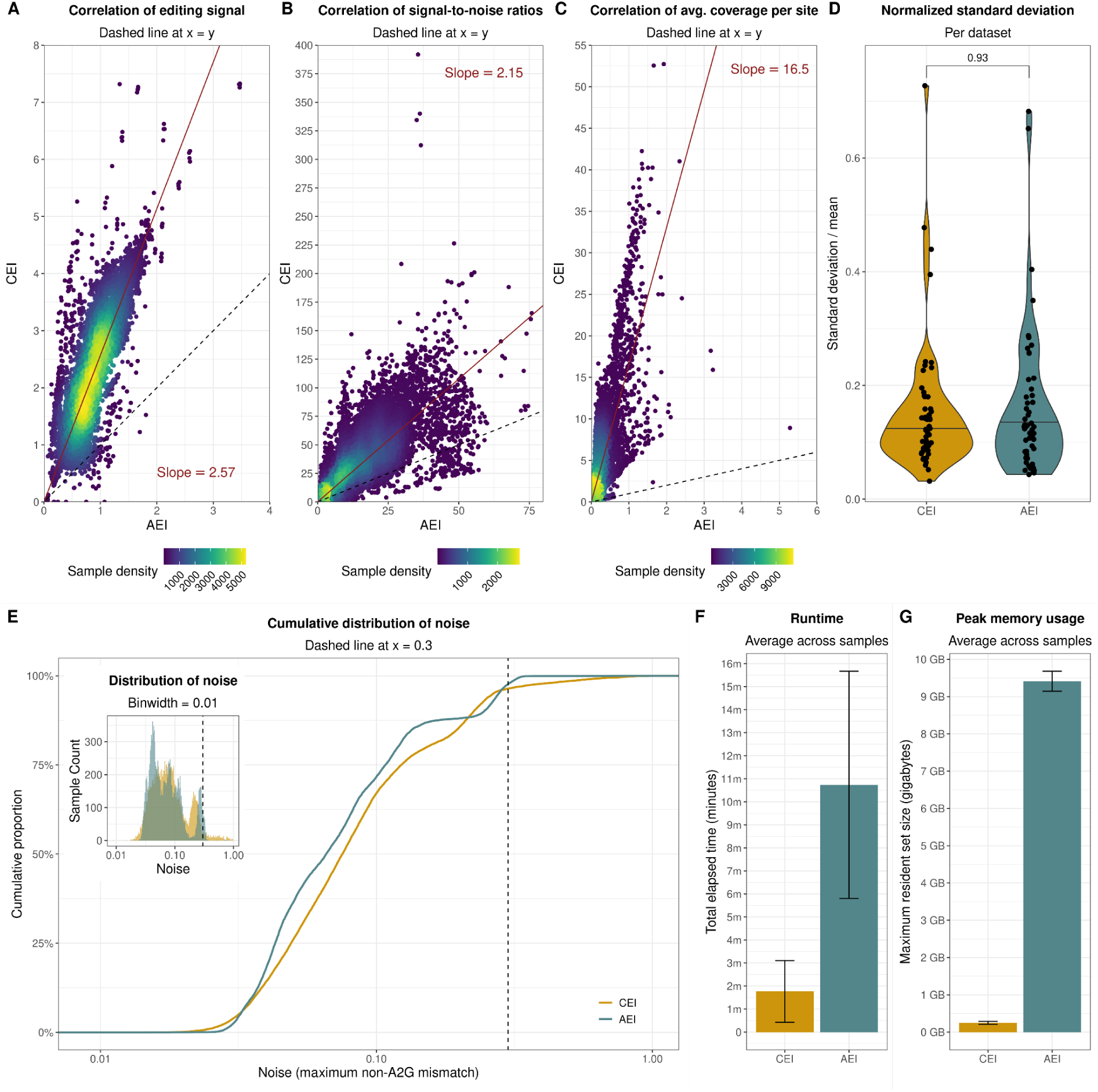
Cytoplasmic editing index demonstrates superior signal, specificity, and efficiency over global editing metrics. *a–c,* Scatter-density plots compare the CEI and AEI in terms of *a*) the editing signal (A-to-G index, *b*) signal-to-noise ratio (SNR), and *c*) mean coverage per editable site. Points are coloured by sample density; dashed black lines mark x = y, and red lines are ordinary-least-squares fits through the origin, giving slopes of 2.57, 2.15 and 16.5, respectively. SNR is the A-to-G index divided by the maximum non-A-to-G mismatch index (four samples with infinite SNR were excluded). Average coverage is computed per editable site (Methods). *d,* Dataset-level variability. Each point is the standard deviation of the index values across all samples within a single dataset normalised by its mean editing level, restricted to samples with noise < 0.3 in both indices (dataset sizes 22–1,299 samples, Supplementary Table S9). No significant difference is found between indices (Wilcoxon signed-rank). *e,* Empirical cumulative distribution of editing noise (maximum non-A-to-G mismatch index per sample). Although the two curves largely overlap, the global index has a slightly larger fraction of samples under the threshold of 0.3 (vertical dashed line), consistent with slightly lower background mismatch variability. Inset: noise density histogram. *f,g,* Computational efficiency. Bars show the mean ± s.d. wall-clock runtime per sample (*f*) and peak resident-set memory per sample (*g*), both recorded with the GNU time utility. Both metrics are substantially lower for the CEI, underscoring its performance advantage.

### The cytoplasmic editing index increases sensitivity to editing alteration in infection

We next compared use of CEI and AEI to detect immune-related editing alterations at the population level across diverse infections. Using our uniformly processed compilation of 15,649 RNA-seq samples spanning more than 20 pathogens and associated controls, we computed CEI and AEI for each sample (Fig. 5a), and searched for infection-associated editing changes (Fig. 5b).

**Figure 5:**
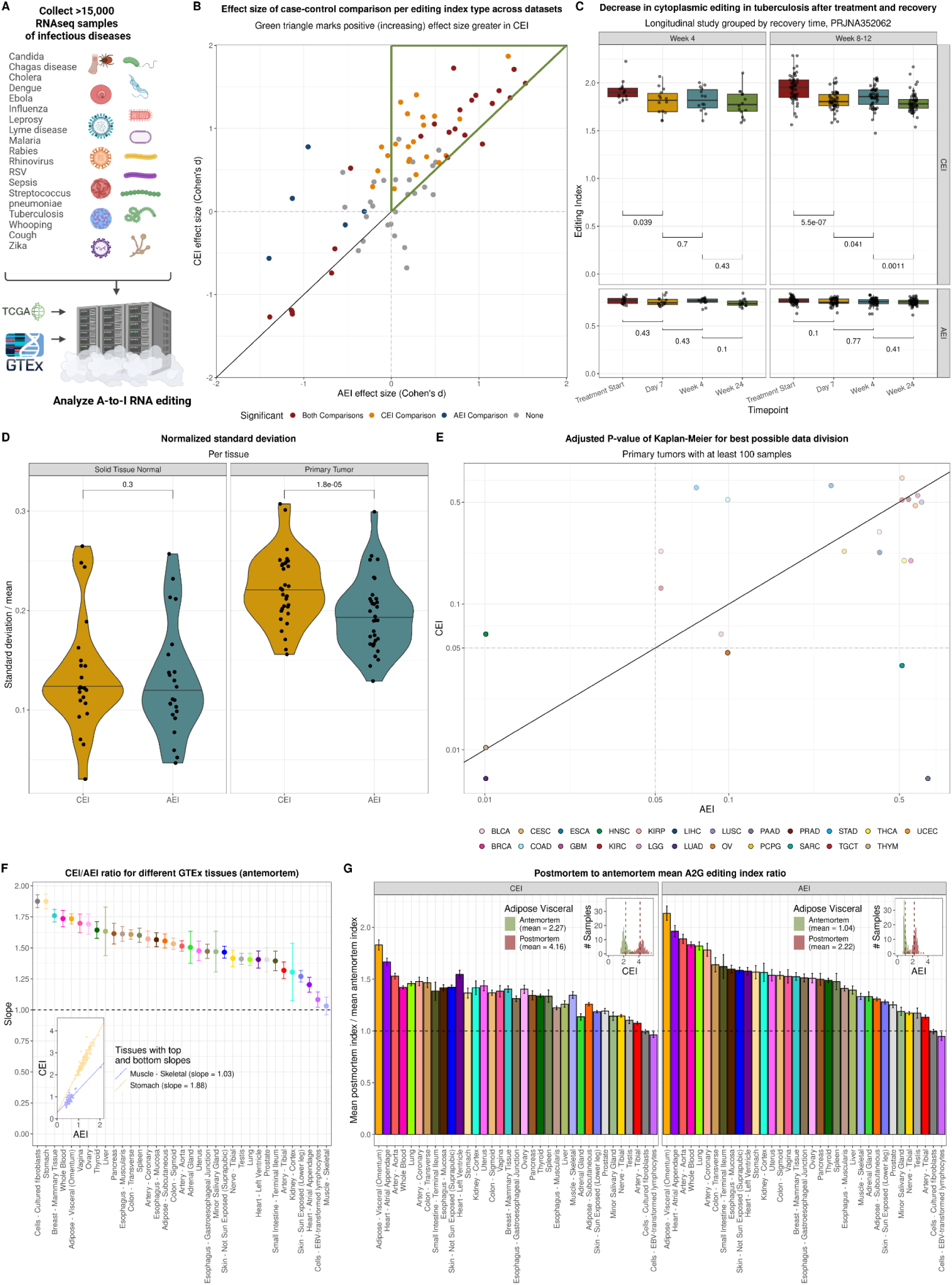
Cytoplasmic editing index enables detection of immune-regulated RNA editing changes. *a,* Overview of RNA-seq sample collection and analysis workflow. A total of 15,649 publicly available RNA-seq samples spanning 55 datasets and over 20 distinct infectious diseases were compiled and analyzed using standardized cloud-based pipelines to quantify the CEI and AEI. *b,* Comparison of effect sizes (Cohen’s d, mean difference divided by pooled standard deviation) for case-control analyses between CEI and AEI across infectious disease datasets (for all case-control comparisons with ≥4 samples per group, ≥10 total samples). The green triangle indicates the region where the CEI identified a greater positive effect size than the AEI, demonstrating enhanced biological sensitivity. Points are colored based on statistical significance detected exclusively by CEI, exclusively by AEI, by both indices, or by neither index (Wilcoxon test, paired samples where applicable, see Methods). Case–control status was assigned from sample metadata and pathogen load annotations (Methods; Supplementary Table S1; Supplementary Table S6). *c,* Longitudinal CEI analysis of tuberculosis treatment and recovery^78^ in human samples. Boxplots show editing index trajectories at several time points after treatment initiation, grouped by recovery time (either week 4 or weeks 8–12). The CEI decreases significantly after recovery, while the AEI shows no significant changes (Wilcoxon rank-sum test, raw p-values denoted under comparisons). *d,* Variability of RNA editing indices across TCGA cancer samples. Each point corresponds to one tissue type, and presents the normalized standard deviation (s.d./mean editing) of the index across samples in primary tumors and normal tissues. Inter-tumor variability of CEI is significantly greater than AEI in primary tumors (Wilcoxon signed-rank test), whereas normal tissues show no significant difference between indices. *e,* Kaplan-Meier survival analysis across TCGA cancer types with ≥100 samples. Adjusted P-values (Benjamini–Hochberg correction) represent the optimal separation of samples into two groups based on the editing index (Methods). The two indices showed complementary patterns of survival association, with CEI performing better in some cancer types and AEI in others. *f,* Linear modeling (CEI ∼ AEI) across antemortem donors of GTEx for tissues with ≥10 samples. Points represent model slopes (± standard error) per tissue, ranked by slope magnitude. Inset shows linear fits for tissues with the highest (stomach) and lowest (muscle skeletal) slopes, highlighting variable tissue-specific relationships between cytoplasmic and global editing indices. g, Postmortem vs. antemortem editing index stability across GTEx tissues with ≥10 samples in both cohorts. Bars indicate postmortem-to-antemortem mean editing ratios for CEI and AEI, highlighting the reduced susceptibility of CEI to postmortem RNA degradation. Insets show distributions of postmortem and antemortem CEI and AEI values in adipose visceral tissue, exemplifying the robustness of CEI against tissue-degradation artifacts.

Across infections, CEI consistently showed larger absolute effect sizes and greater sensitivity, achieving statistical significance in 46 comparisons, compared to only 28 significant changes in AEI (Fig. 5b). This advantage was especially pronounced for editing increases during active infection, consistent with CEI’s sensitivity to ADAR1p150 upregulation, and was most evident for viral infections and respiratory conditions (Supplementary Fig. S7). In a longitudinal tuberculosis cohort^78^, CEI captured a progressive decrease in editing after treatment and recovery (Fig. 5c), whereas AEI showed no significant changes, highlighting CEI’s capacity to track dynamic, immune-driven regulation of RNA editing.

### Improved detection of RNA editing variability in cancer and across normal tissues using the CEI

We next examined whether the enhanced sensitivity of CEI extended beyond infectious diseases. We first looked at variability and patient stratification in cancer samples and controls from the TCGA dataset. CEI captured greater editing variability than AEI across primary tumors but not normal tissues, underscoring its capacity to detect heterogeneity in tumor-associated editing changes (Fig. 5d). Furthermore, Kaplan–Meier survival analyses showed that CEI and AEI provided better patient stratification in different cancer types across TCGA, highlighting complementary strengths and underscoring the clinical value of using both metrics (Fig. 5e). CEI showed stronger signal and lower noise relative to AEI in TCGA tumors, and the indices differed in the cancer types they most strongly distinguished, reflecting complementary strengths (Supplementary Fig. S9).

To assess the robustness of CEI across tissues and donor cohorts, we analyzed normal samples from both antemortem (living samples: organ donor, surgical) and postmortem donors, taken from the GTEx^77^ dataset. The editing index is affected by postmortem processes, such as hypoxia, cellular metabolism and apoptosis^39,60^, making this distinction critical for evaluating tissue-specific measurements. The correlation between CEI and AEI varied substantially across tissues, revealing that different tissues exhibit specific immune-relevant editing landscapes (Fig. 5f; Supplementary Fig. S8). As a notable exception, cortex-derived brain tissues uniquely exhibited CEI values lower than the AEI ones, potentially reflecting distinct editing dynamics driven by high expression of ADAR2 and ADAR3 in this neural tissue^47,79^ (Supplementary Fig. S8). Importantly, CEI was less susceptible to postmortem changes (Fig. 5g; Supplementary Fig. S8). These findings underscore the stability of CEI across donor cohorts and support its use for quantifying RNA editing in both high-quality and partially degraded tissues.

## Discussion

Quantifying immunologically relevant A-to-I RNA editing represents a critical challenge in understanding innate immune regulation, as existing metrics fail to capture the biological specificity required for mechanistic insights. The Cytoplasmic Editing Index (CEI) provides a biologically informed complement to existing RNA editing metrics by targeting the subset of editing events most likely to influence cytoplasmic dsRNA sensing.

The qualitative difference between CEI and global editing metrics is manifested in CEI’s selective ADAR1p150 dependence. Furthermore, the significant depletion of inverted *Alu* clusters relative to random expectations provides an evolutionary validation for CEI’s targeting strategy, indicating a strong selective pressure against dsRNA-forming arrangements, consistent with previous findings^66^. This evolutionary constraint, shown here for human and mouse, confirms that the repeats targeted by CEI targets are enriched in functionally critical editing substrates.

Our scalable, cost-effective computational pipeline delivers substantial technical advantages, and is particularly suitable for large-scale or heterogeneous datasets. Per-sample processing costs are reduced to ∼$0.10 while maintaining reproducibility through containerized, cloud-native workflows. This proved crucial for our analysis of >25,000 samples across diverse contexts, demonstrating that CEI consistently outperformed AEI with higher editing signal, improved signal-to-noise ratio and higher coverage per editable site, without increased variability. These advantages translated into enhanced biological detection across infectious diseases, complementary patient stratification for cancer contexts, and reduced susceptibility to postmortem artifacts.

Importantly, our implementation of the CEI is highly accessible and enables widespread adoption. The user is required to supply just a list of SRA accession codes, and all steps from data download to index calculation are fully automated.

As a targeted index prioritizing specificity over completeness, CEI focuses on *Alu* sequences in RefSeq 3′UTRs and may not capture alternative dsRNA structures such as non-*Alu* repeats, *Alu* elements that are expressed in long 3’UTRs that not represented in RefSeq sequences, or even polymorphic sequences that are not part of the reference genome. Future extensions could extend the genomic scope and integrate RNA secondary structure mapping to identify additional immunostimulatory regions.

CEI’s enhanced sensitivity suggests a possible clinical utility for monitoring immune status in infectious diseases and autoimmune conditions where ADAR1 dysfunction drives pathology. The improved cancer stratification hints at potential prognostic applications, though prospective validation would be required. By targeting functionally meaningful substrates rather than aggregate editing, CEI provides a powerful analytical framework for dissecting how A-to-I editing shapes immune regulation in health and disease, complementing existing global metrics.

## Methods

### Terminology

**Alu element:** A single *Alu* element >200 bp as defined in the RepeatMasker database. It may or may not overlap other *Alu* elements.

***Alu* cluster:** The set of all *Alu* elements located within a 3’UTR of a curated RefSeq transcripts. *Alu* elements that overlap the 3’UTR partially and exceed beyond its limits are trimmed to include only the subsequence overlapping with the 3’UTR. Two distinct 3’UTR isoforms may define two distinct *Alu* clusters.

**Inverted *Alu* cluster:** An *Alu* cluster containing at least one element of each orientation.

**Tandem *Alu* cluster:** An *Alu* cluster that is not an inverted *Alu* cluster, where all *Alu* elements have the same orientation.

### Infectious Disease Data Collection and Annotation

Fifty-five publicly available infection-related RNA-seq datasets, comprising 15,741 RNA-seq samples, were obtained from the NCBI Sequence Read Archive (SRA)^80^. Datasets were annotated via SRA metadata, with additional curation from GEO^81^ SOFT files when metadata were incomplete. The datasets are listed in Supplementary Table S1. Accession numbers, metadata, and per-sample annotations are summarized in Supplementary Table S2.

Each study was classified as case-control or alternative design (e.g., longitudinal, treatment). For longitudinal datasets, two time-points were selected as case and control based on study specifications, as detailed in Table S1. The time-point with the highest pathogen load was chosen as the case group whenever possible, unless a different selection criterion was noted in the primary publication.

For multi-strain pathogens, strains were analyzed separately only if each strain group contained at least four samples. Otherwise, strains were combined into a single group. Datasets containing multiple libraries from the same donor were treated as independent observations; no donor-level random-effect correction was performed. Pathogen classification by clinical manifestation and pathogen type is listed in Supplementary Table S5. Case-control comparisons are detailed in Supplementary Table S6.

Of 15,741 samples in this cohort, 92 failed to run due to various issues, most commonly because of mismatched paired-end lengths (e.g., 50 bp + 150 bp) or missing FASTQ files. The failed samples and a detailed breakdown of failure reasons are provided in Supplementary Table S4.

### ADAR1 Perturbation Data Collection

In addition, we analyzed 147 publicly available RNA-seq samples from datasets profiling genetic or cytokine perturbations of RNA editing regulators in human and mouse along with cell fractions^10,70–76^. These datasets included experimental conditions such as ADAR knockdown or reconstitution, as well as interferon stimulation time courses along with sequencing of cell fractionation protocol. Accession numbers, metadata, and sample groupings for these experiments are provided in Supplementary Table S3.

### GTEx Data Collection and Annotation

Raw RNA-seq data for 9,602 samples from the Genotype-Tissue Expression v10 (GTEx)^77^ project were obtained via dbGaP^82^ (phs000424). Only normal samples sequenced with paired-end, unstranded, short-read protocols with available FASTQ files via SRA were included. Surgical and organ-donor samples were classified as antemortem (tissues collected from living donors), whereas postmortem samples originated from deceased donors. Accession numbers, metadata, and sample group assignments for all analyzed GTEx samples are provided in Supplementary Table S8.

### TCGA Data Collection and Analysis

Aligned RNA-seq data from The Cancer Genome Atlas^83^ (TCGA) were obtained from the Genomic Data Commons (GDC)^84^ via dbGaP (phs000424). We excluded multi-mapped and unmapped reads, to retain only uniquely mapped primary alignments compatible with our reference annotations, using the command samtools view -h -b -q 255 -F 2308 -f 2. Of the initial 11,083 samples, 452 samples were excluded due to the absence of reads after filtering. Accession numbers, metadata, filtering outcomes, and sample group assignments for all analyzed TCGA samples are provided in Supplementary Table S7. Detailed workflow adaptations for TCGA data are documented and available on GitHub.

### Cloud-Based Computational Pipeline and Data Processing

All computational analyses used our cloud-optimized Nextflow pipeline engineered for cost-efficient processing of high-throughput RNA-seq data. The workflow runs on Google Cloud Platform (GCP) Batch and Amazon Web Services (AWS) Fargate, but is portable to other providers owing to its modular design. All steps are executed using version-pinned

Docker images to ensure reproducibility across analyses. Stage-specific resource allocation reduced average end-to-end cost to ∼$0.10 per sample (Fig. 1c).

**Initialization:** a one-time initialization step retrieves all reference annotations and generates index files required for expression quantification, alignment and RNA quantification. The resources are then deposited in a bucket of the user’s choice. This step is done once per computing platform and organism.

**Per-sample processing:** this includes per-sample tasks — read retrieval, quality control, expression quantification, alignment to the genome, and RNA-editing quantification. For each sample, these steps are executed sequentially in discrete containers on resource-optimized virtual machines. Samples are processed in parallel, enabling thousands of concurrent workflows and simplifying quota management and monitoring. The workflow accepts a list of SRA accession IDs (and an NGC file, for dbGaP data) and produces sample-level outputs ready for downstream analysis.

**Retrieving all reference annotations:** reference annotations and expression files are downloaded from UCSC Genome Browser^85^ (http://hgdownload.soe.ucsc.edu/goldenPath; https://hgdownload.soe.ucsc.edu/gbdb) canonical data sources and processed using standard genomics tools (awk, bedtools^86^ and bigBedToBed^85^). Specific resource URLs are detailed in GitHub.

**Generation of index files required for expression quantification:** Salmon^87^ (version 1.10.2) index was generated with RefSeq^85^ (latest available version as of early 2024) as the reference transcriptome. The entire genome was used as the decoy sequence as per Salmon’s comprehensive index generation protocol.

**Generation of index files required for alignment:** for each read-length group we built a length-matched STAR^88^ index (--sjdbOverhang = R, where R ɛ {50, 75, 100, 125, 150}).

**Generation of index files required for RNA editing quantification:** for each regions file, we built the appropriate JSD index using the RNAEditingIndexer. This index is used during the editing quantification step to avoid per-sample re-generation and reduce resource usage.

**Resource deposit:** resources are transferred to a bucket supplied by the user via AWS CLI or gcloud CLI, according to the cloud provider.

**Read retrieval and quality control:** raw sequencing reads were retrieved using NCBI SRA Toolkit (v3.0.7), and preprocessed with fastp^89^ (v0.23.4) for quality filtering, with non-default options -q 25 -u 20 -e 30 --dont_eval_duplication -G -A. For each dataset (library), we extracted X, the average read length per mate as annotated in SRA. We defined L as the element in the set of standard lengths (L ɛ {50, 75, 100, 125, 150}), that is closest to X. If X ≥ L-3, reads shorter than L-3 bp were discarded using the flag --length_required L-3. Reads longer than L were trimmed to this size from the 3’ using --max_len1 L. Else (X < L-3), reads shorter than X-3 bp were discarded using the flag --length_required X-3, and reads longer than X were trimmed to this size from the 3’ using --max_len1 X. In both cases, the alignment index for this data was set to L. In some cases, SRA annotations were found to be inconsistent with read data and were manually curated. All read lengths decisions for both X and L are detailed in Supplementary Table S2 at sample level. Studies containing multiple read-length groups (e.g., 50 bp and 100 bp libraries) were split, and each group of specific read-length was processed independently. GTEx samples were processed using X = L = 75bp as per their documented sequencing protocol.

**Expression quantification:** transcript expression levels were quantified using Salmon^87^ (version 1.10.2) with automatic library detection (-l A) and transcript-to-gene mapping (-g) using a relevant resource file.

**Alignment to the genome:** unique alignments were generated with STAR^88^ (v2.7.10b), with the relevant index and the following parameters:

--alignSJoverhangMin 8 --alignIntronMax 1000000 --alignMatesGapMax 600000

--outFilterMultimapNmax 1 --outFilterMatchNminOverLread 0.95. The index length was chosen to match preprocessed read lengths as previously detailed. For indices robustness testing, we selected 1,000 random samples and realigned them with the non-AEI-optimal default STAR parameter --outFilterMatchNminOverLread 0.66 while keeping other flags unchanged. Comparison of noise is presented in Supplementary Fig. S5b.

RNA-editing quantification: RNA editing levels within *Alu* regions (AEI, CEI, tandem index) were quantified from STAR alignments using RNAEditingIndexer^39^ (v1.0) with RefSeq curated data as reference. Common variants from dbSNP151^90^ (hg38) for human samples or dbSNP142 (mm10) for mouse samples were filtered to exclude known polymorphisms from the analysis. RNA editing levels of known sites of human^91^ and mouse^92^ are quantified via pipeline using a custom-built program based on samtools^93^ mpileup file format.

Resource usage monitoring: runtime and peak memory usage were measured using the GNU time utility and aggregated across samples to benchmark computational efficiency. Results are presented in Fig. 2f and Fig. 2g.

The complete Nextflow workflow, pipeline initialization and documentation are available at https://github.com/a2iEditing/CEI. All known sites and CEI annotations, including pre-computed inverted *Alu* cluster definitions for human and mouse reference genomes (hg38 and mm10 accordingly), are provided.

### Cytoplasmic Index Regions

The cytoplasmic editing index is calculated over inverted *Alu* clusters. These were identified by scanning 3′UTR exons for ≥2 *Alu* elements (each exon separately), each spanning *>*200 bp, with at least one *Alu* element in each of the two strand orientations. *Alu* elements ≤200 bp were excluded, even when positioned inside a candidate cluster. For mouse regions, B1 and B2 families were analyzed separately, with clusters considered inverted when at least one element within a family showed inverted orientation relative to other elements in the same family. Mouse repeats were required to span *>*120 bp.

Elements extending beyond annotated 3′UTR boundaries were trimmed to retain only the sequence overlapping the 3’UTR. Transcriptomic annotations were obtained from UCSC Genome Browser^85^ (hg38 and mm10 RefSeq Curated, February 2021). Adjacent or overlapping *Alu* elements were merged with pybedtools^94^ (v0.9.1, wrapping bedtools^86^ v2.30). The lists of clusters and associated elements are available in Supplementary Table S15, Supplementary Table S16 and the project’s GitHub repository.

### Tandem Index Regions

Tandem index regions are *Alu* elements that belong to tandem *Alu* clusters, do not belong to any inverted *Alu* cluster, and in addition do not have an inverted *Alu* element of any length (even ≤200 bp) occurring anywhere in the same 3′UTR, to ensure structural independence from potential dsRNA-forming configurations. The same transcript annotations (hg38 and mm10 RefSeq Curated, February 2021) and 3’UTR-trimming rules were applied. Adjacent elements were merged with pybedtools^94^ (v0.9.1; bedtools v2.30) using the same parameters as for inverted clusters. For mouse samples, tandem regions were defined separately for B1 and B2 families. The resulting tandem set represents non-dsRNA-forming repetitive element clusters and serves as a control for evaluating orientation-specific editing activity.

### RNA Editing Index Calculations

RNA editing levels were quantified using RNAEditingIndexer^39^ (v1.0). Two editing indices were computed for all samples, with a third index calculated for specific analyses:

Global editing index (AEI): quantified editing levels across all annotated *Alu* elements genome-wide using default hg38 settings for human samples or mm10 settings for mouse samples (B1 and B2 elements).

Cytoplasmic editing index (CEI): quantified editing within inverted *Alu* clusters (or inverted B1/B2 elements for mouse) as defined above, using the -rb flag. This metric restricts analysis to cytoplasm-localized, dsRNA-forming clusters and serves as the primary biologically-informed alternative to the AEI.

Tandem index: calculated for control analyses (Fig. 3) using non-inverted 3′UTR *Alu* elements via the -rb flag as a structural control.

All indices used mismatch profiles across specified regions, masking common SNPs (dbSNP151 for hg38; dbSNP142 for mm10). RNAEditingIndexer incorporates gene expression estimates to guide strand assignment and transcript disambiguation during editing quantification (for algorithmic details, see Ref. ^39^).

### Inverted Cluster Count Model

To compute the expected ratio of the numbers of inverted and tandem clusters, we noted that of all 2ᴺ possible allocations of orientation to elements in a cluster with N *Alu* elements, only two configurations result in a tandem cluster — either all forward or all reverse. Assuming each element is independently oriented, with equal probability for forward or reverse orientation, one expects an inverted-to-tandem ratio:

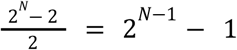

### Calculation of average coverage per editable site

Mean coverage per editable site is calculated as the total read depth at all A and T positions in the reference sequence, divided by the number of such positions. We excluded contributions from mismatches other than A-to-G and T-to-C. Both forward strand A positions and reverse strand T positions are included since A-to-I editing can occur on either strand depending on transcript orientation.

### Statistical Analysis of Infectious Diseases Cohort

Case–control comparisons in infection datasets were assessed with Wilcoxon rank-sum tests on noise-filtered data (maximum non-A-to-G mismatch < 0.3); Wilcoxon signed-rank tests were applied when study design allowed. Comparisons were considered only when filtered case and control groups included ≥ 4 samples each and ≥ 10 samples in total. If a dataset included multiple tissues or read lengths, each tissue and read length were compared separately. Technical replicates were merged after noise filtering by averaging over editing measures across all replicates. For paired comparisons, unpaired samples (due to experimental design or filtering) were excluded.

### Calculation of Standard Deviation for Ratio of Means

Standard deviations for ratios of group means were calculated using error propagation. For each group, the sample mean and standard deviation were computed using the unbiased estimator (denominator n-1). We estimated the coefficient of variation as 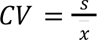 for each group.

For the ratio 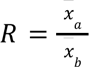 the variance was calculated using the delta method, assuming independence between groups. The standard deviation of the ratio is:

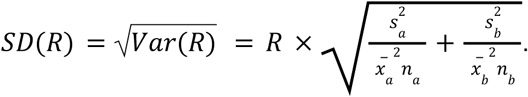

### Linear Modeling and Comparative Analysis of GTEx Cohort

To compare the two indices using linear modeling, we first filtered out noisy samples (maximum non-A-to-G mismatch index ≥0.3 for either AEI or CEI). Donor replicates within tissues were merged by averaging over editing measures across all replicates. The filtered and merged cohort included 9,097 samples.

Ordinary least squares regression (CEI ∼ AEI, no constant term) was applied to each tissue type with ≥10 samples, to find the slopes. To compare antemortem and postmortem cohorts, we repeated the analysis for each cohort separately, requiring ≥10 samples in both cohorts.

### Survival Analysis and Comparative Analysis of TCGA Cohort

Following noise filtering and sample merging as described above, the TCGA cohort included 10,488 samples.

For each TCGA cancer type with ≥100 samples, we split the data based on index values to perform survival analysis. Samples were ranked by editing index value and divided into high and low groups using nine different thresholds at 10% increments (10th percentile, 20th percentile, …, 90th percentile, see Supplementary Table S13). For each threshold, Kaplan–Meier^95^ survival analysis was performed in R v4.5.0 using the survival package (v3.8-3) (`survfit()`) and log-rank testing (`survdiff()`). P-values were adjusted for multiple testing (for all cancer types and thresholds, CEI and AEI separately) using the Benjamini-Hochberg procedure to control the false discovery rate, and the threshold yielding the lowest log-rank P-value was chosen. This approach ensures fair comparison between indices by allowing each to be evaluated at its optimal discrimination threshold, while controlling for the multiple comparisons inherent in the threshold optimization procedure.

### Statistics

All statistical tests presented in the article are two-sided.

### Reproducibility and Data Availability

All code, pipeline definitions and configuration files are publicly available at https://github.com/a2iEditing/CEI. Custom Docker images are archived at https://hub.docker.com/u/levanonlab. Raw RNA-seq data were downloaded from the NCBI Sequence Read Archive (SRA)^80^; accession numbers are provided in Supplementary Tables S2 and S3. Controlled-access data from TCGA^83^ (dbGaP accession phs000178) and GTEx^77^ (dbGaP accession phs000424) were obtained under approved data-use agreements; accession numbers are provided in Supplementary Tables S7 and S8.

All computational steps were executed in version-pinned Docker containers orchestrated by Nextflow^96^ (v25.03.1-edge) to ensure end-to-end reproducibility. Statistical analyses were conducted in R v4.5.0 using tidyr v1.3.1, dplyr v1.1.4, purrr v1.0.4, data.table v1.17.6 and stringr v1.5.1. Publication-ready figures were generated in R using ggplot2 v3.5.2, cowplot v1.1.3 and patchwork v1.3.1. Venn diagrams were generated with DeepVenn^97^. Illustrative figures were created with <u>BioRender.com</u>.

Data generated or analysed during this study are included in this published article and its Supplementary Information files.

## Acknowledgements

We thank the following members of the Levanon research group for their contributions to initial dataset search and curation: Aviya Zimmermann, David Gorelik, Zohar Rossenwasser-Maymon, Michelle Eidelman, Miriam Karmon, Moshe Perez, Netanel Landesman, Kobi Shapira, Rivka Bouhnik-Cohen, and Renana B. Drummer. We are grateful to Yamin Cohen for compiling the clinical manifestation categories. We acknowledge the numerous research groups, including the GTEx Consortium, who deposited RNA-seq data in public repositories and made this large-scale comparative analysis possible. The results shown here are in whole or part based upon data generated by the TCGA Research Network: https://www.cancer.gov/tcga.

## Author contributions

EYL and EE conceived the study and supervised the work. EYL, EE and RCF developed the CEI methodology. IT designed and implemented the cloud platform workflow and conducted the sample analysis. Resource initialization and data analysis were performed by RCF. IT and RCF annotated infectious disease. HK collected and curated infectious disease datasets and contributed to the initial analyses. SHR designed and implemented the known-sites editing quantification program and developed the Docker container. EYL, EE and RCF wrote the manuscript with feedback from IT.

## Supplementary materials

### Supplementary figures

**Supplementary Figure S1:**
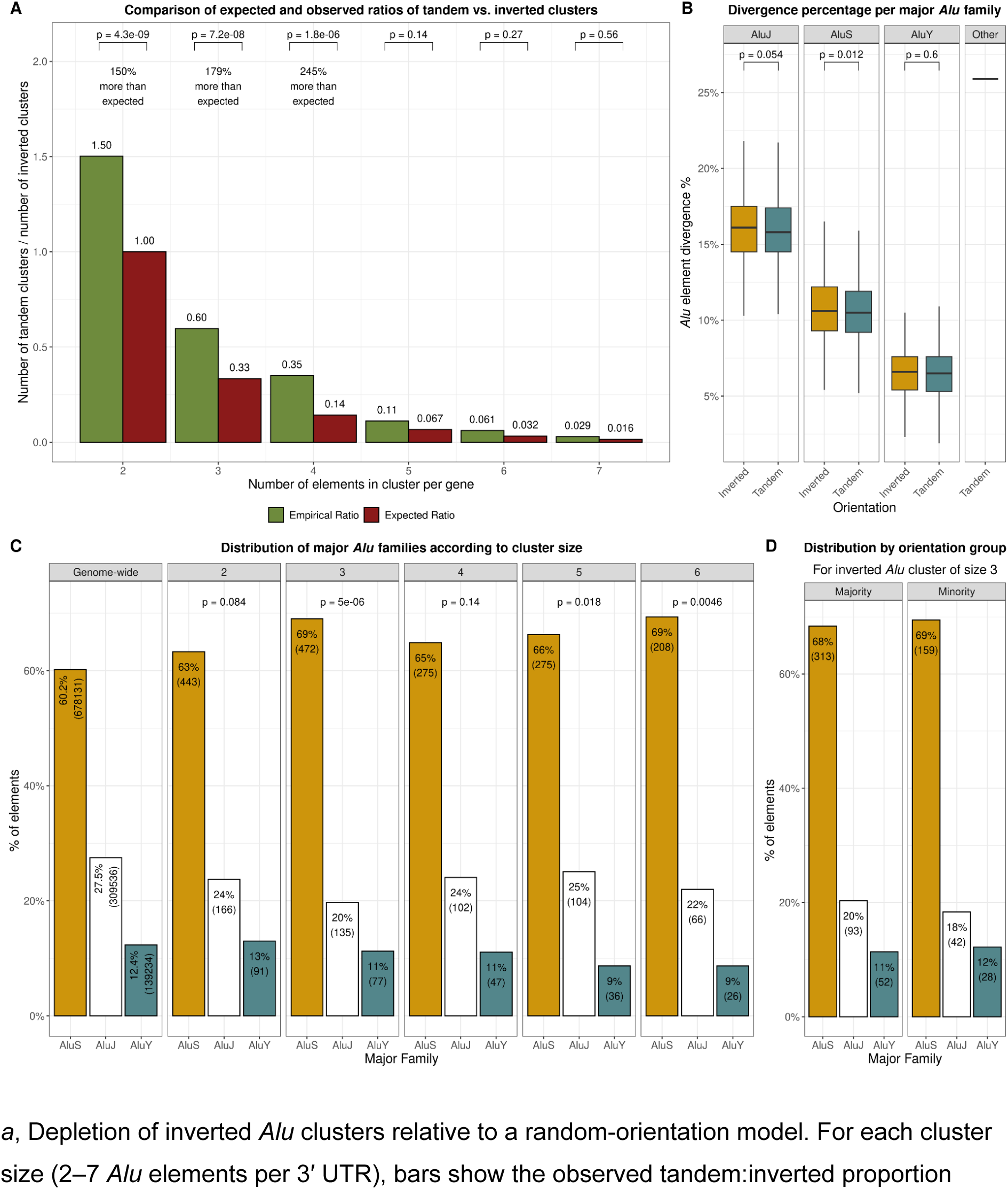
Additional properties of inverted and tandem *Alu* clusters in human 3′ UTRs. *a*, Depletion of inverted *Alu* clusters relative to a random-orientation model. For each cluster size (2–7 *Alu* elements per 3′ UTR), bars show the observed tandem:inverted proportion (green) and the proportion expected under a random null model, assigning random, equally probable and uncorrelated orientations (red; see Methods for derivation). Numbers above bars indicate the fold excess of tandem clusters over expectation. Inverted *Alu* clusters are significantly depleted for clusters at sizes 2–4 (χ² test, uncorrected p-value), consistent with selective pressure against dsRNA-forming arrangements. *b,* Divergence from consensus for *Alu* elements in inverted versus tandem clusters, stratified by major family (AluJ, AluS, AluY). No significant change was found for AluJ and AluY families, and only a modest difference in divergence of AluS elements (Wilcoxon test, no multiple testing correction for the three tests). *c*, Composition of major *Alu* families within inverted *Alu* clusters by cluster size. For each size class (2–6) and for the genome-wide baseline, bars give the percentage (and count) of AluS, AluJ and AluY elements. AluS is enriched across all sizes (χ² test, no multiple testing correction for the five tests). *d*, Family composition of majority- versus minority-orientation elements within inverted clusters of size 3. Here, “majority” refers to the two *Alu* elements that share the same strand within a size-3 inverted cluster, whereas ‘minority’ denotes the single element on the opposite strand. Bars show the percentage and count of each family in the two orientation subsets, revealing similar family distributions.

**Supplementary Figure S2:**
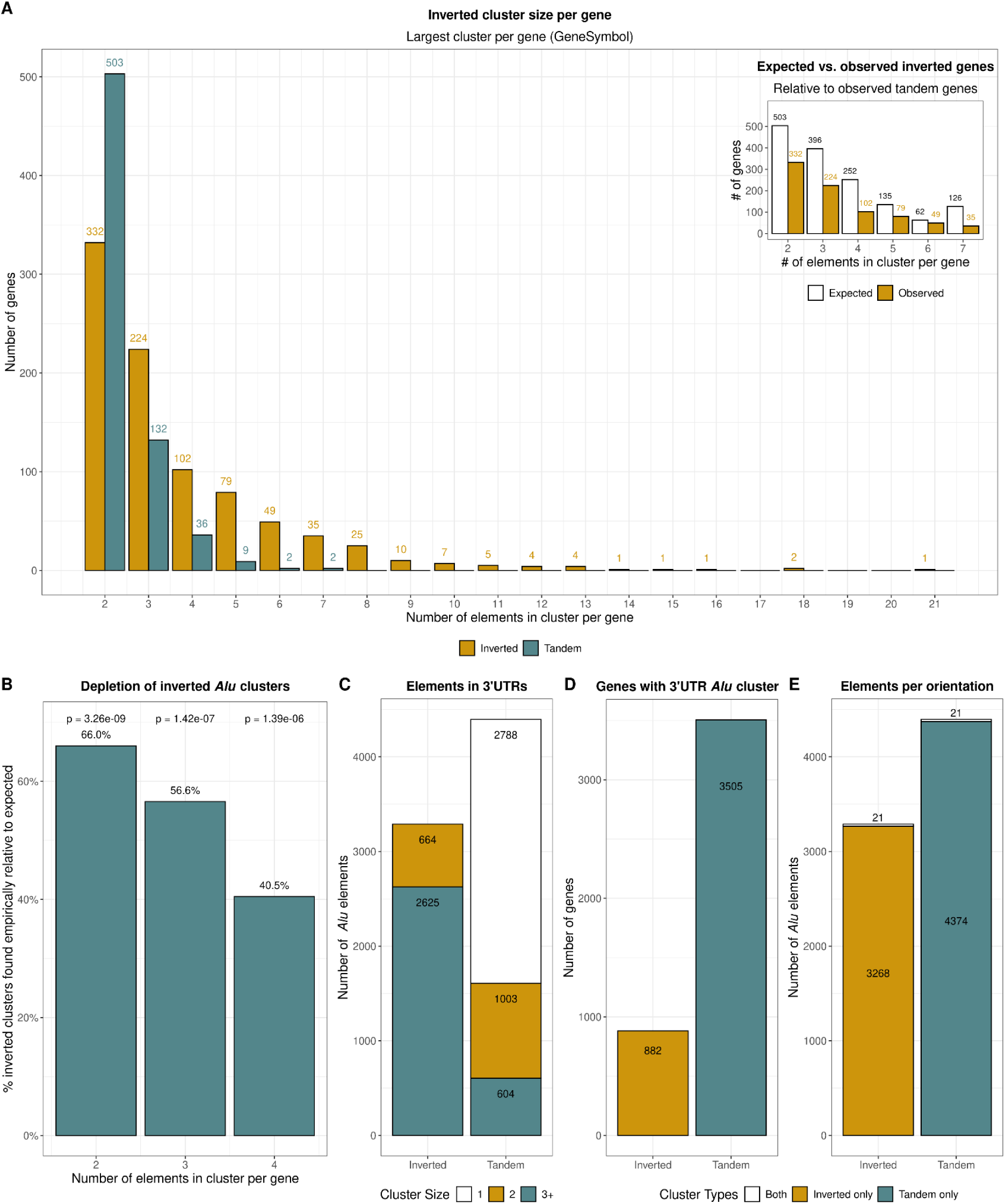
Orientation architecture of *Alu* clusters after longest-isoform selection. Panels *a–d* repeat the analyses of Fig. 2 *b–d,f* after restricting each gene to its longest 3′ UTR (see Fig. 2 legend for methodological details); the isoform collapse removes gene double-counting, so that each gene is classified as either inverted or tandem. The resulting size distributions (*a*), depletion of inverted *Alu* clusters relative to random-orientation model (*b*), element counts by orientation class (*c*), and gene-level prevalence (*d*) all recapitulate the trends observed in Fig. 2, confirming that orientation biases and the rarity of inverted, dsRNA-forming *Alu* clusters are robust to isoform choice. *e,* Element-level overlap of orientation classes. Bars show the number of *Alu* elements unique to the inverted class, unique to the tandem class, or found in both classes (i.e., inverted in one gene and tandem in another). The overlap highlights individual elements that adopt different structural roles in different gene contexts under the longest-isoform definition.

**Supplementary Figure S3:**
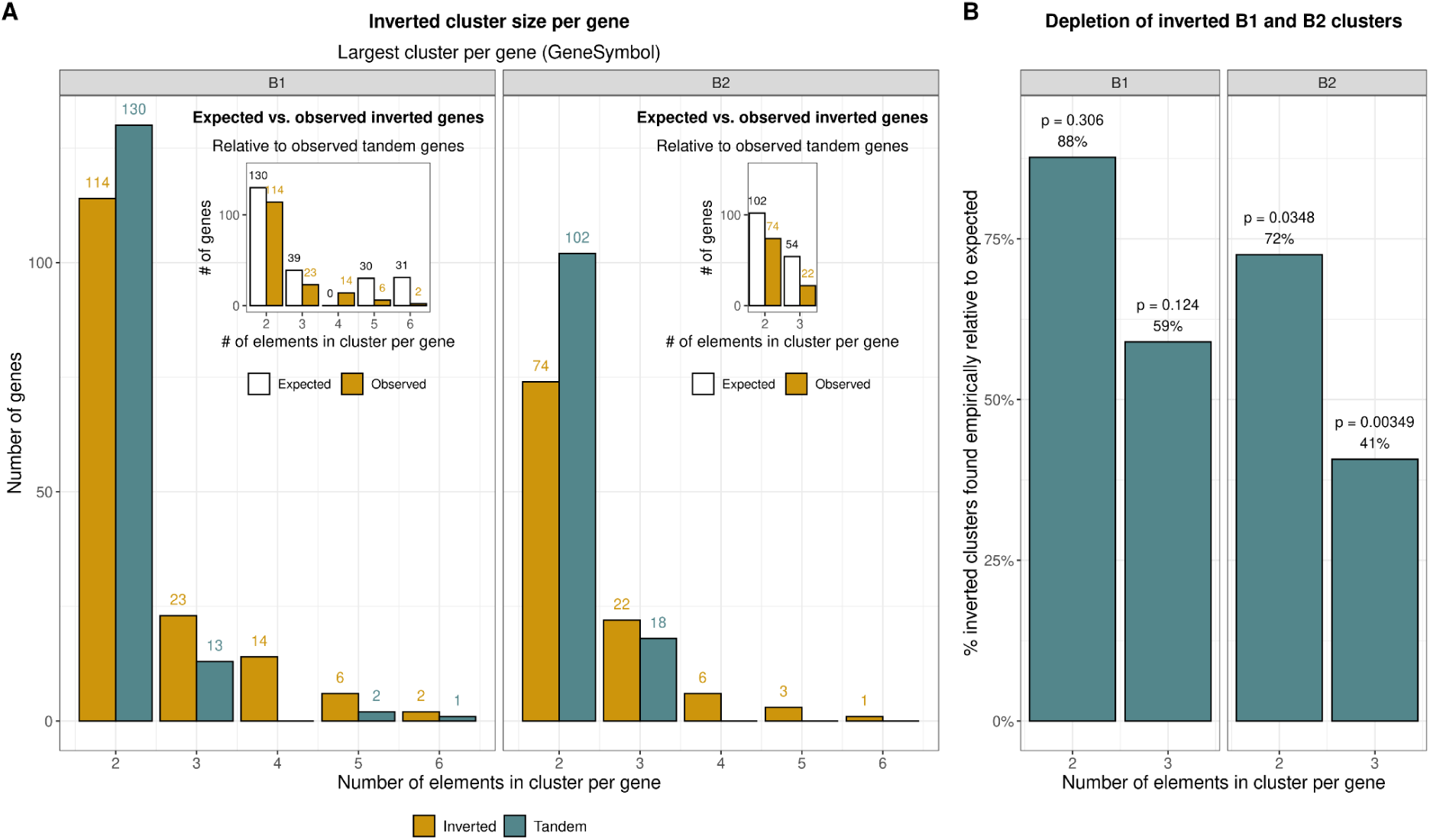
Structural constraint on inverted versus tandem B1 and B2 clusters in mouse 3′UTRs. *a,* Distribution of the largest B1 and B2 cluster per gene (defined as the 3’ UTR with the greatest number of elements), stratified by orientation. Numeric labels above each bar give the number of genes in each category. Inverted clusters extend to larger sizes than tandem clusters, reflecting the greater combinatorial space of mixed orientations, yet remain rarer than expected (see b). Inset panels compare the observed number of genes harbouring inverted B1 clusters (left inset) or inverted B2 clusters (right inset) with the counts expected under the random-orientation model (see Methods). *b,* Depletion of inverted clusters relative to a random-orientation model. For cluster sizes 2–3, bars plot the observed proportion of inverted clusters divided by the proportion expected when orientations are assigned at equal probability (Methods). Values < 100% indicate depletion; B2 shows significant depletion (*χ*² test, uncorrected p-value).

**Supplementary Figure S4:**
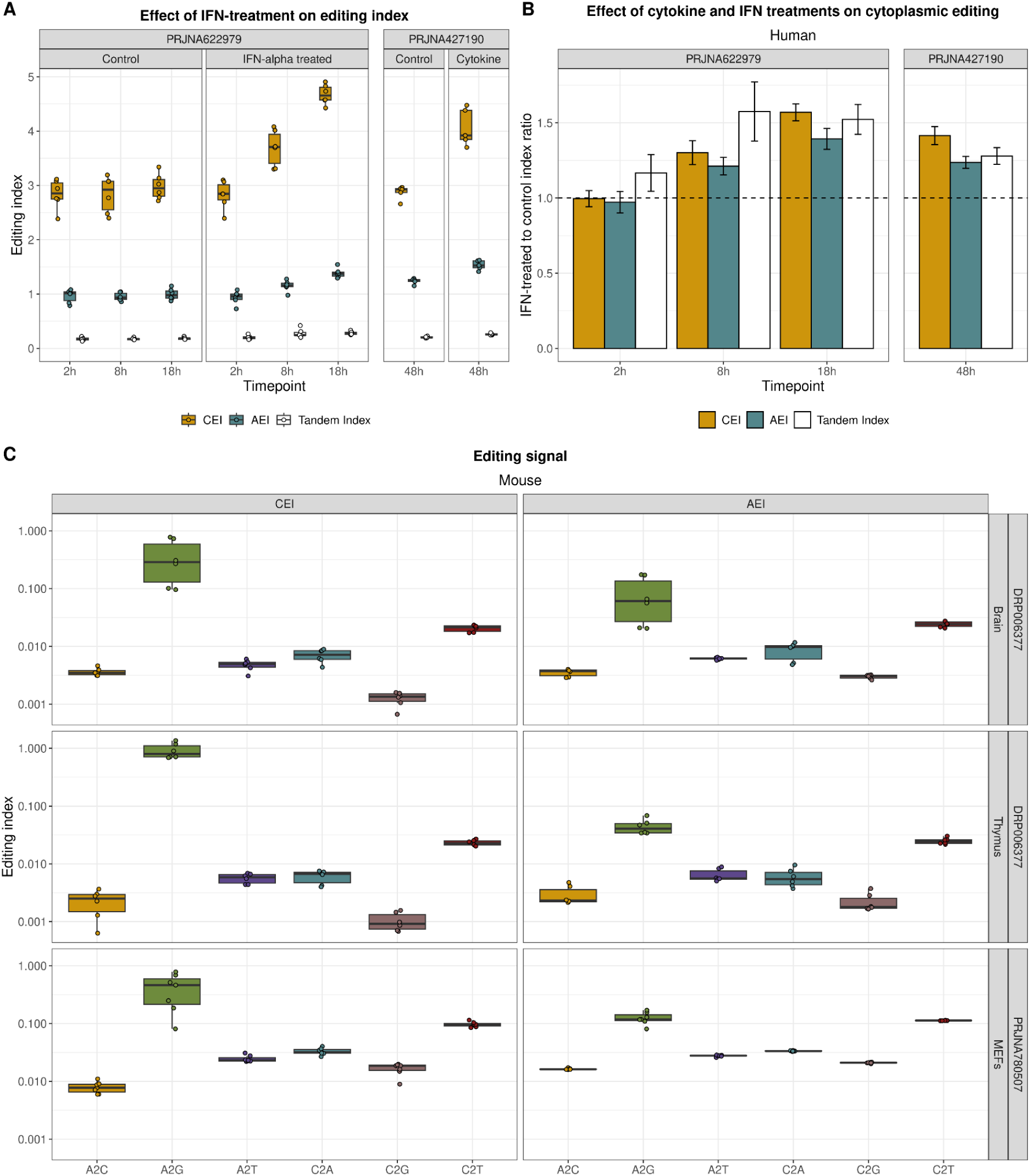
Cytokine-induced cytoplasmic editing in human islets and mismatch specificity in mouse datasets. *a*, Temporal dynamics of cytokine-induced editing in primary human islets. Two additional independent datasets (IFN-α, left^71^; mixed cytokines, right^73^) demonstrate the cytoplasmic editing index rising earlier and more substantially than the global *Alu* editing index. The tandem index remains consistently low and largely unaffected by cytokine treatment. *b*, Fold-induction (ratio of mean cytokine-treated to mean control editing indices) for datasets shown in panel *a*. The CEI consistently exhibits greater dynamic induction compared to the AEI. Apparent fold-changes in the tandem index do not reflect biologically meaningful induction, as the absolute editing levels remain near baseline (panel *a*). *c*, Mismatch specificity in mouse datasets. Editing index distributions for each mismatch type shown for mouse brain, thymus, and MEFs datasets. While the A-to-G editing signal is consistently and markedly elevated in the CEI relative to the AEI, all non-A-to-G mismatches (noise) remain comparable between the two indices, confirming the CEI’s enhanced sensitivity to genuine editing events without elevating technical noise.

**Supplementary Figure S5:**
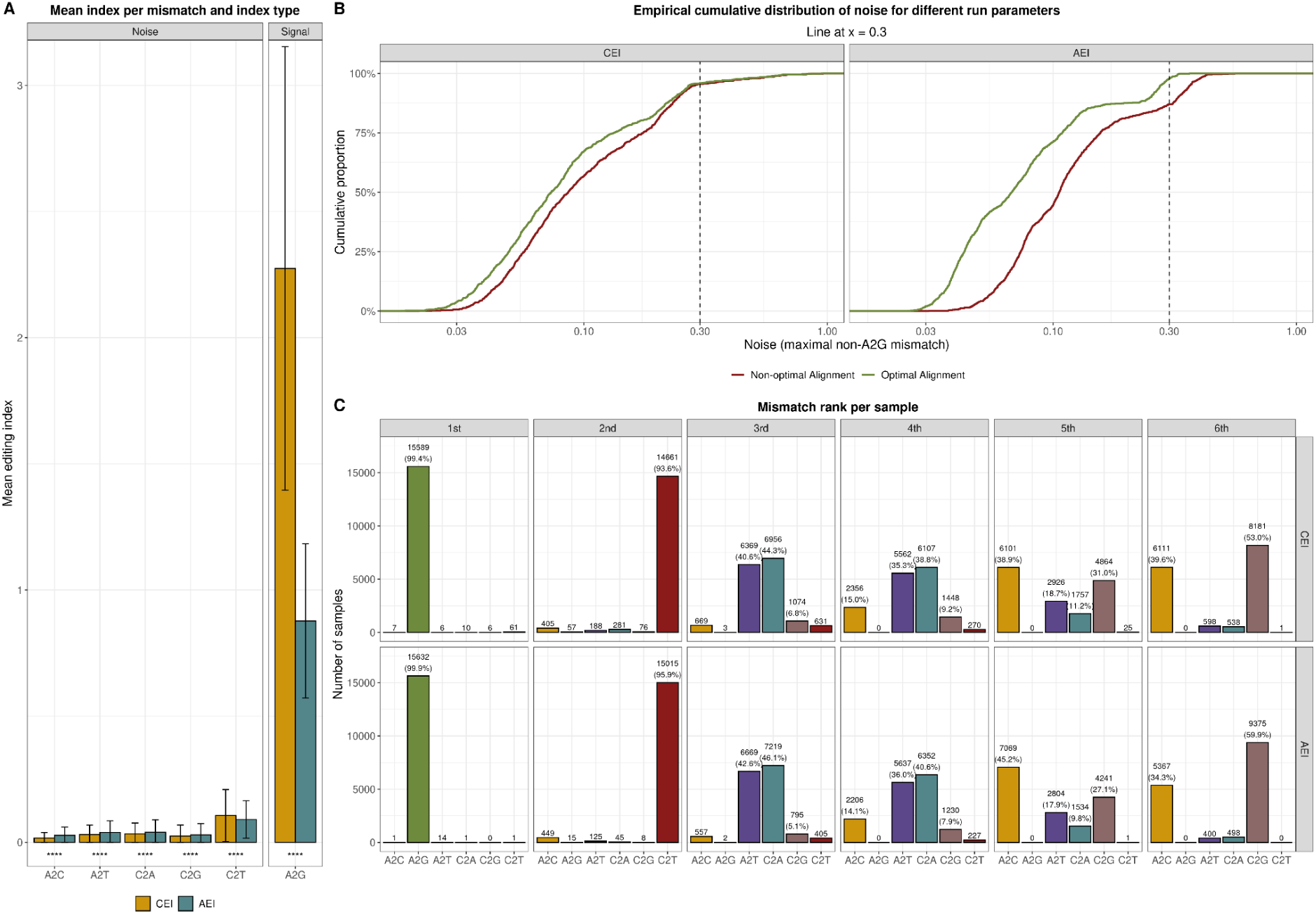
Mismatch characteristics of the CEI versus the AEI. *a*, Mean editing index (± s.d.) for each nucleotide transition. Compared with the global index, the cytoplasmic index markedly enriches the A-to-G signal and reduces all non-A-to-G mismatches except C-to-T, which is modestly higher — consistent with the C-to-U being the most prevalent mutation and SNP. All pairwise differences are significant (two-sided Wilcoxon, **** *P* < 2.2 × 10⁻¹⁶). *b,* Robustness to alignment parameters. A random set of 1,000 samples was re-aligned with STAR using the AEI-optimised parameter set reported in the original study and, for comparison, the STAR default (--outFilterMatchNminOverLread 0.95 vs. 0.66; see Methods for full command lines). Empirical CDFs plot editing noise (maximum non-A-to-G mismatch per sample). The cytoplasmic index curve is less sensitive to alignment quality. The dashed line at 0.30 marks the noise threshold used downstream. *c,* Per-sample ranking of mismatch types (dense ranking of ties). Bars show the number of samples (percentages shown only where > 5%) in which each mismatch ranks 1st–6th for the cytoplasmic (upper row) and global (lower row) indices. A-to-G transitions rank first in over 99% of samples. C-to-T transitions dominate the second rank (over 93% of samples), while other mismatches remain rare at higher ranks. These patterns confirm that genuine A-to-I editing events dominate the mismatch landscape, with C-to-T changes providing a consistent biological background, likely reflecting the biological prevalence of C-to-U deamination (e.g., spontaneous CpG deamination or APOBEC activity).

**Supplementary Figure S6:**
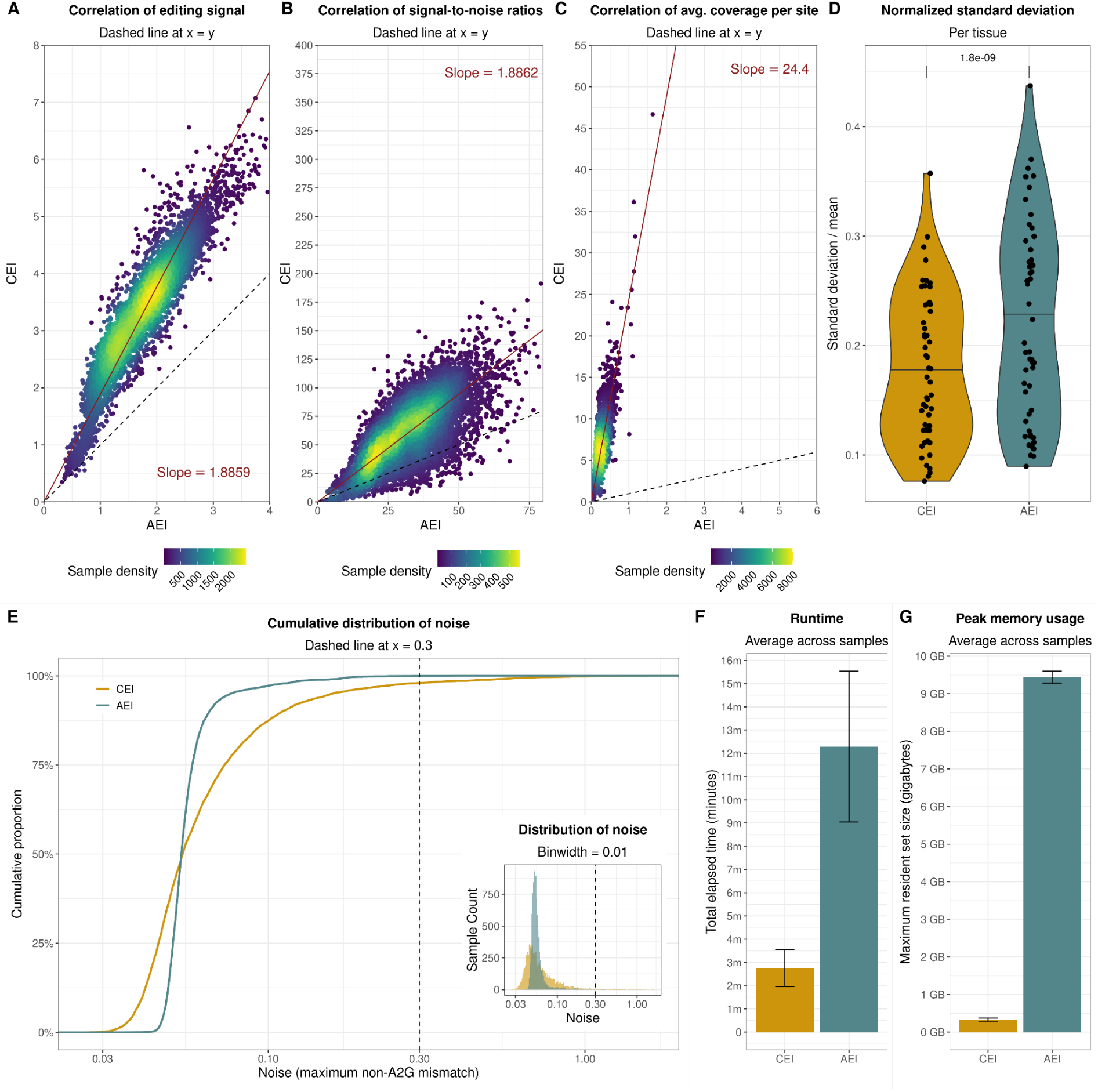
Performance of the cytoplasmic editing index in the GTEx cohort. This figure repeats the analyses of Fig. 4 for GTEx; see the Fig. 4 legend for metric definitions and plotting conventions. Key GTEx-specific results are summarised below. *a–c,* Scatter-density plots comparing the cytoplasmic editing index (CEI) with the global *Alu* editing index (AEI) for editing signal (*a*), signal-to-noise ratio (*b*) and mean coverage per editable site (*c*). Ordinary-least-squares fits through the origin give slopes of 1.89, 1.89 and 24.4, respectively, confirming that the CEI retains stronger signal, higher specificity and deeper coverage in GTEx. *d,* Tissue-level variability. Each point is the standard deviation of one GTEx tissue (53 tissues) normalised by its mean editing level, using only samples with noise < 0.30 in both indices. Variability is significantly lower for the CEI (Wilcoxon signed-rank test). *e,* Empirical cumulative distribution of editing noise. The AEI curve lies slightly above the CEI curve at the 0.30 threshold (vertical dashed line), indicating a marginally larger fraction of low-noise samples for the AEI in this cohort. Inset, noise-density histogram. *f,g,* Computational efficiency remains substantially lower for the CEI, underscoring the CEI’s performance advantage across large uniform cohorts.

**Supplementary Figure S7:**
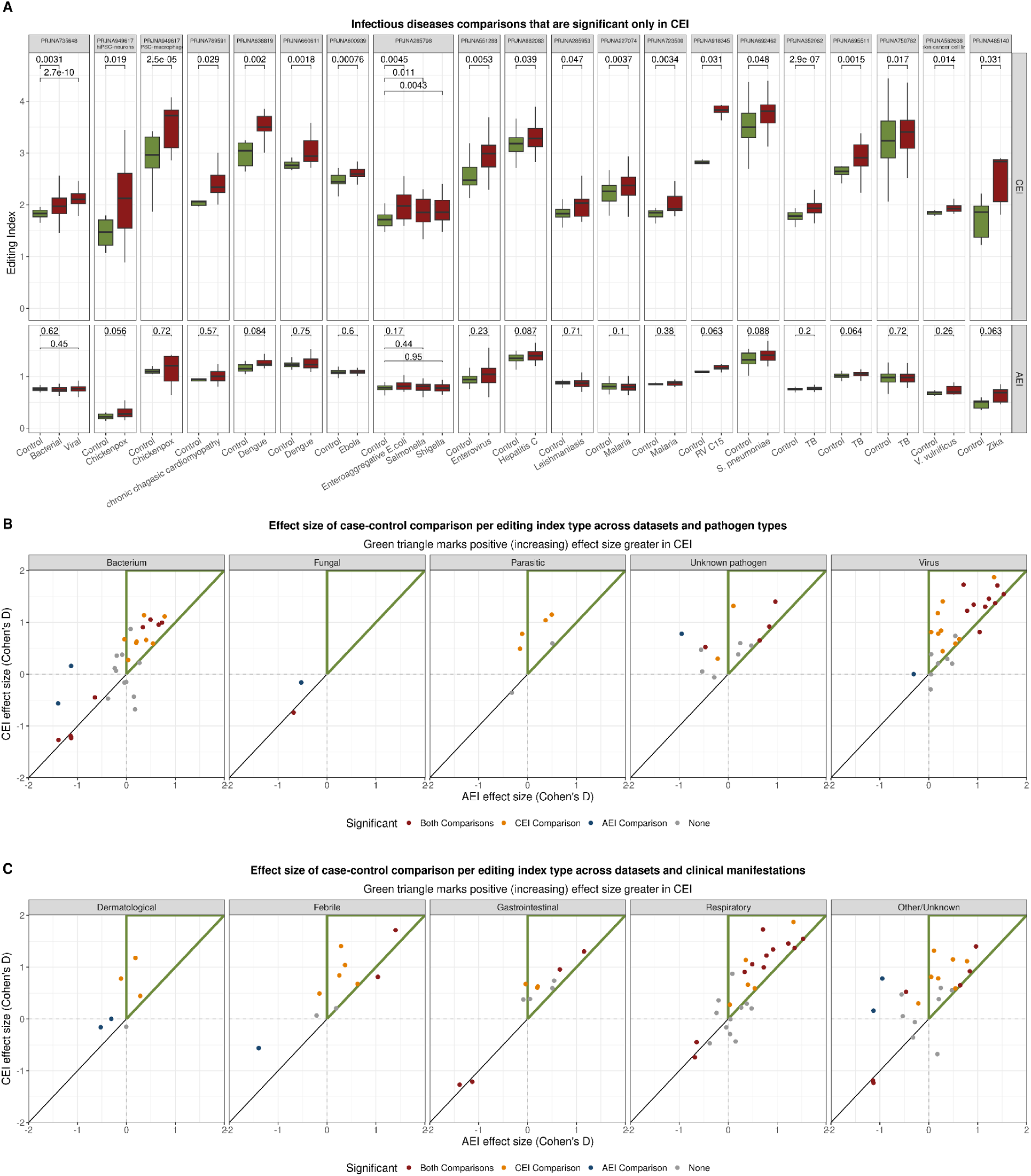
Cytoplasmic editing index captures biologically relevant immune-related editing signals across infectious disease contexts. *a*, Case-control comparisons that show a statistically significant difference in CEI but not in AEI, as shown in Fig. 5b, spanning bacterial, viral, and parasitic infections (all comparisons with maximum non-A-to-G mismatch < 0.3, ≥4 samples per group, ≥10 total samples). Statistical significance determined by Wilcoxon test (raw p-values). *b,c*, Effect-size (Cohen’s D) comparisons stratified by pathogen type (b) and clinical manifestation (c). Points above the diagonal line (within the green triangular region) indicate greater positive effect sizes detected by the CEI relative to AEI. The advantage of CEI is most pronounced in viral infections and respiratory conditions. Points are colored based on statistical significance detected exclusively by CEI, exclusively by AEI, by both indices, or by neither index (Wilcoxon test, paired samples where applicable, see Methods).

**Supplementary Figure S8:**
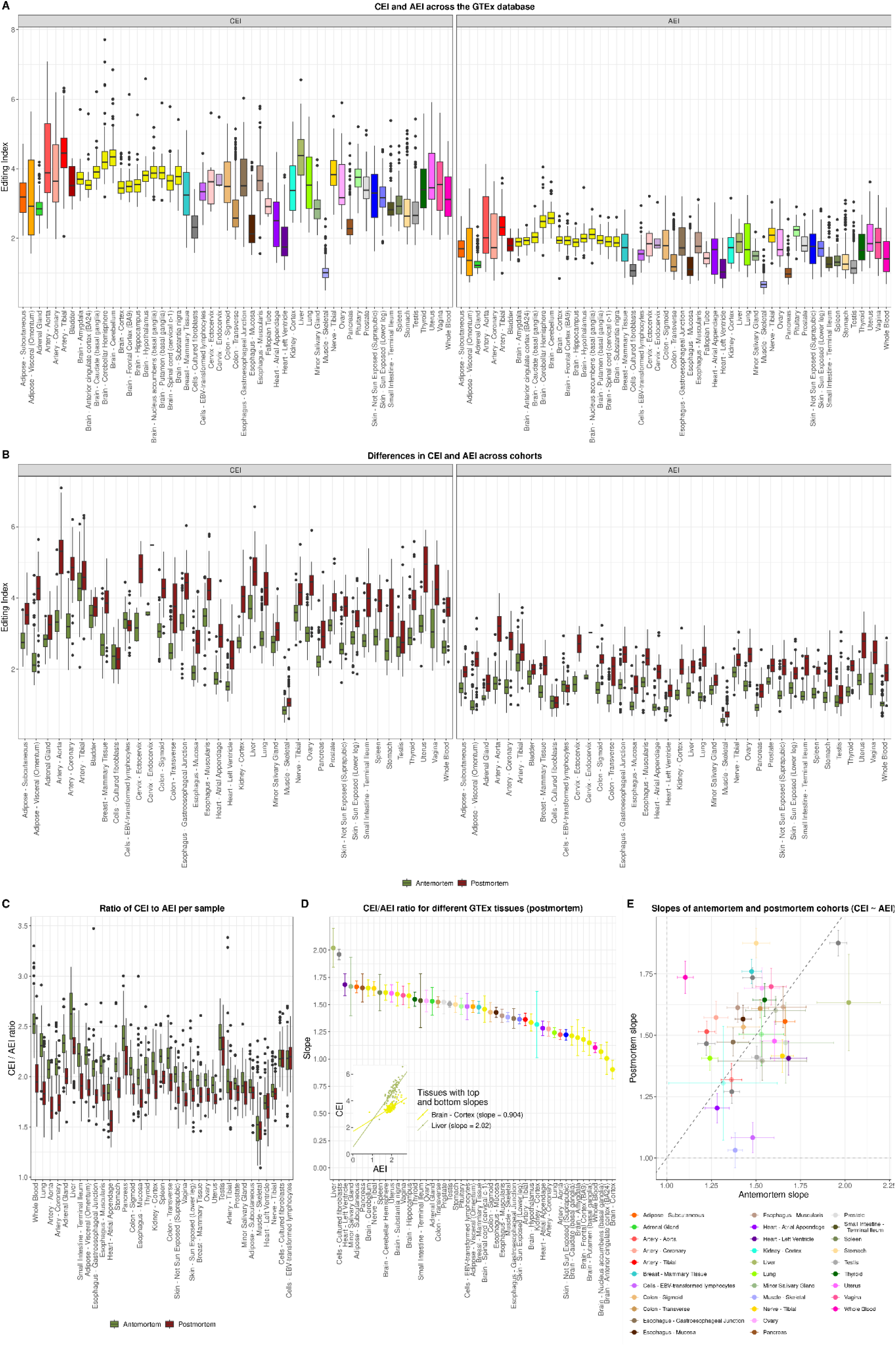
Comparison of cytoplasmic and global RNA editing relationships across GTEx tissues and cohorts. *a*, Distribution of editing indices across GTEx tissues. Boxplots show CEI and AEI values for all tissues, demonstrating tissue-specific variation in both indices and their relative magnitudes. *b*, Editing index distributions stratified by cohort and tissue. Boxplots compare CEI and AEI values between antemortem and postmortem samples across tissues with both cohorts, revealing higher values in postmortem samples. *c*, Distribution of sample-level CEI/AEI ratios, split by antemortem and postmortem donor cohorts. Higher ratios indicate an increased relative contribution of cytoplasmic editing. Only GTEx tissue samples with ≥10 samples in both cohorts are shown. *d*, Linear modeling (CEI ∼ AEI) across postmortem GTEx tissue samples with ≥10 samples. Points represent model slopes (± standard error) per tissue, ranked by slope magnitude. This postmortem analysis complements the antemortem analysis shown in Fig. 5f, highlighting tissue-specific variation in the relationship between cytoplasmic and global editing across cohorts, including distinct patterns within brain tissues. *e*, Comparison of CEI ∼ AEI model slopes between antemortem and postmortem donor cohorts. Each point represents a tissue with ≥10 samples in both cohorts. Points above the diagonal line indicate tissues where the CEI-AEI relationship is stronger in postmortem samples, while points below indicate stronger relationships in antemortem samples. This analysis evaluates consistency in tissue-specific editing relationships between cohorts.

**Supplementary Figure S9:**
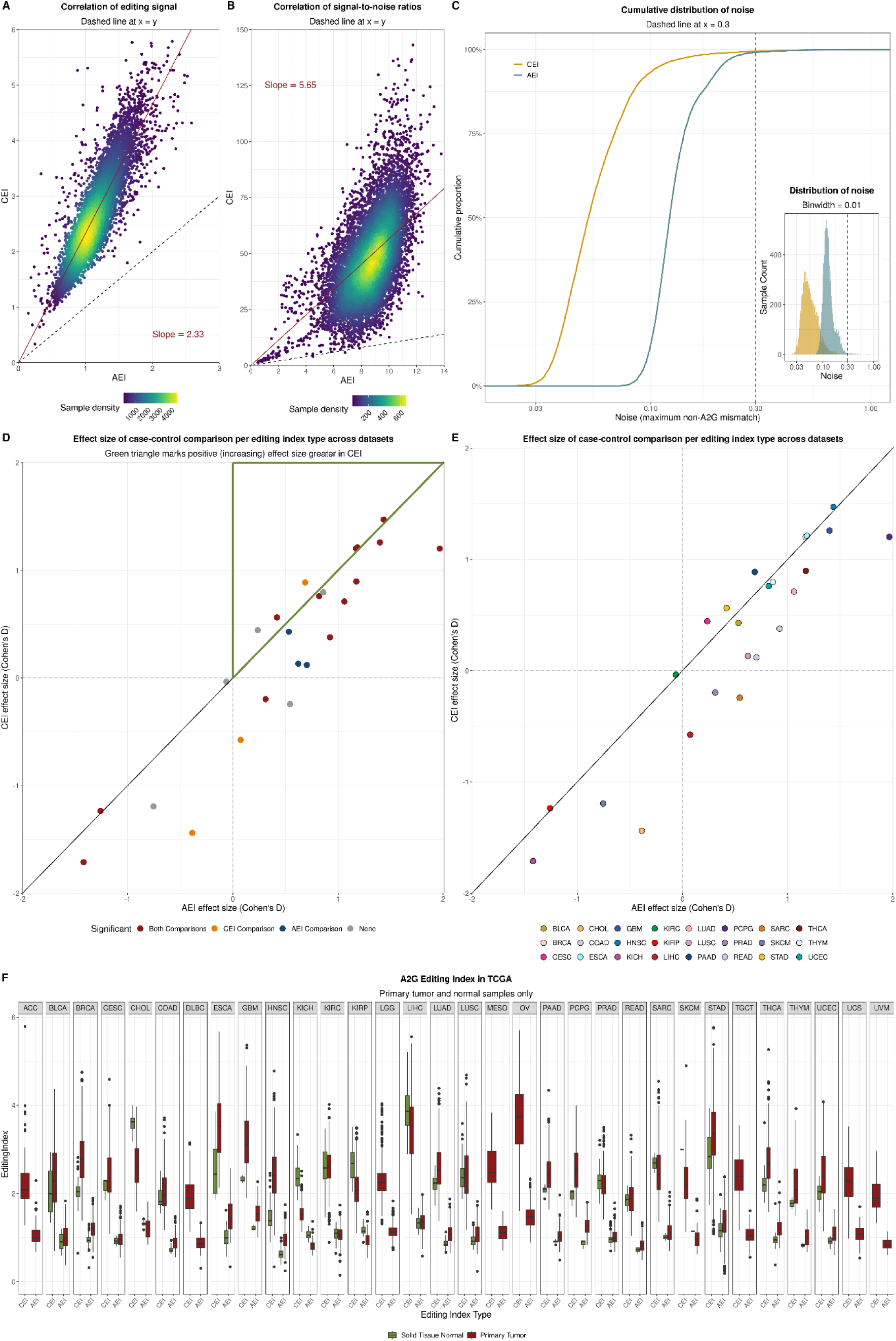
Signal quality metrics and cytoplasmic editing patterns in TCGA cancer samples. *a,b,* Scatter-density plots compare the CEI and AEI for editing signal (A-to-G index, *a*) and signal-to-noise ratio (SNR, *b*). Points are coloured by sample density; dashed black lines mark x = y, and red lines are ordinary-least-squares fits through the origin. SNR is the A-to-G index divided by the maximum non-A-to-G mismatch index. These metrics validate CEI measurements in cancer cohorts, demonstrating that the cytoplasmic index delivers stronger editing signal and higher specificity than the global index. *c,* Empirical cumulative distribution of editing noise (maximum non-A-to-G mismatch index per sample). The CEI shows slightly lower background noise levels compared to the AEI, consistent with the CEI’s greater robustness to alignment parameters (Supplementary Fig. S5) when applied to GDC alignments. Inset: noise density histogram. *d,e,* Comparison of effect sizes (Cohen’s d, mean difference divided by pooled standard deviation) for tumor-normal analyses between CEI and AEI across TCGA cancer types (for comparisons with ≥10 samples per group, noise threshold <0.3 for both indices). Points represent cancer types, colored by statistical significance (*d*) or tissue of origin (*e*). Green triangle (d) denotes comparisons where the CEI identified a greater positive effect size than the AEI (Wilcoxon test, paired samples where applicable). *f,* Cytoplasmic editing index distributions across all solid tumors and normal samples. Boxplots compare CEI values between tumor and normal tissues, providing an overview of editing patterns across the cancer cohort.

## Description of Supplementary tables

Table S1: Infectious-disease dataset annotation

Per-dataset metadata for all infectious-disease studies analysed, including study design, pathogen, disease indication and technical descriptors.

Table S2: Infectious-disease sample annotation

Per-sample metadata for the infectious-disease cohort, detailing original SRA/GEO columns and annotation, curated case–control annotations, study design, read length, replicate status, library type, strand specificity and tissue.

Table S3: ADAR perturbation sample classification

Sample classification for ADAR1-perturbation and subcellular fractionation experiments used in Fig. 3.

Table S4: Failed samples

List of infectious-disease samples excluded from analysis, with the specific failure reason for each.

Table S5: Infectious disease taxonomy

Mapping of infectious-disease studies to pathogen type and clinical presentation (source for Supplementary Fig. S7).

Table S6: Infectious-disease case–control contrasts

All case-control comparisons used to compute effect-size differences between editing indices (Fig. 5B; Supplementary Fig. S7).

Table S7: TCGA sample classification

Classification of TCGA samples retained in the study and those removed during filtering (see Methods).

Table S8: GTEx sample classification

Classification of GTEx samples included in the analysis.

Table S9: Infectious-disease datasets: normalised s.d.

Per-dataset summary statistics (mean and normalised standard deviation) for both editing indices (source for Fig. 4D).

Table S10: ADAR perturbation datasets editing metrics

Editing statistics for ADAR1-perturbation and fractionation datasets (Fig. 3; Supplementary Fig. S4).

Table S11: Infectious-disease editing metrics

Editing, coverage and runtime statistics for all infectious-disease samples (Fig. 4, Fig. 5, Supplementary Fig. S5, Supplementary Fig. S7).

Table S12: TCGA editing metrics

Editing statistics for TCGA samples (Fig. 5; Supplementary Fig. S9).

Table S13: Raw survival-analysis output

Kaplan-Meier log-rank results for all cancer types and all high/low index dichotomies (Fig. 5E).

Table S14: GTEx editing metrics

Editing, coverage and runtime statistics for GTEx samples (Fig. 5; Supplementary Fig. S6; Supplementary Fig. S8).

Table S15: Human *Alu* elements in 3’UTRs

Full list of human inverted and tandem *Alu* elements > 200 bp, with the 3′ UTR regions in which they occur (Fig. 2; Supplementary Fig. S1). An element may appear in multiple 3′ UTRs of the same or different genes and may be classified as inverted or tandem in different isoforms.

Table S16: Mouse B1 and B2 elements in 3’UTRs

Full list of mouse inverted and tandem B1 and B2 elements > 120 bp, with the 3′ UTR regions in which they occur (Fig. 2e,g; Supplementary Fig. S3). An element may appear in multiple 3′ UTRs of the same or different genes and may be classified as inverted or tandem in different isoforms.

## References

1. Bass, B. L. RNA editing by adenosine deaminases that act on RNA. Annual review of biochemistry 71, 817–846 (2002).

2. Savva, Y. A., Rieder, L. E. & Reenan, R. A. The ADAR protein family. Genome biology 13, 252 (2012).

3. Nishikura, K. A-to-I editing of coding and non-coding RNAs by ADARs. Nature reviews. Molecular cell biology 17, 83–96 (2016).

4. Schlee, M. & Hartmann, G. Discriminating self from non-self in nucleic acid sensing. Nat Rev Immunol 16, 566–580 (2016).

5. Chen, Y. G. & Hur, S. Cellular origins of dsRNA, their recognition and consequences. Nat Rev Mol Cell Biol 23, 286–301 (2022).

6. Hu, S.-B. et al. ADAR1p150 prevents MDA5 and PKR activation via distinct mechanisms to avert fatal autoinflammation. Molecular Cell 83, 3869–3884.e7 (2023).

7. Mannion, N. M. et al. The RNA-Editing Enzyme ADAR1 Controls Innate Immune Responses to RNA. Cell Reports 9, 1482–1494 (2014).

8. Liddicoat, B. J. et al. RNA editing by ADAR1 prevents MDA5 sensing of endogenous dsRNA as nonself. Science 349, 1115–1120 (2015).

9. Pestal, K. et al. Isoforms of RNA-Editing Enzyme ADAR1 Independently Control Nucleic Acid Sensor MDA5-Driven Autoimmunity and Multi-organ Development. Immunity 43, (2015).

10. Chung, H. et al. Human ADAR1 Prevents Endogenous RNA from Triggering Translational Shutdown. Cell 172, 811–824.e14 (2018).

11. Miyamura, Y. et al. Mutations of the RNA-specific adenosine deaminase gene (DSRAD) are involved in dyschromatosis symmetrica hereditaria. American journal of human genetics 73, 693–699 (2003).

12. Nejentsev, S., Walker, N., Riches, D., Egholm, M. & Todd, J. A. Rare variants of IFIH1, a gene implicated in antiviral responses, protect against type 1 diabetes. Science 324, 387–389 (2009).

13. Rice, G. I. et al. Mutations in ADAR1 cause Aicardi-Goutières syndrome associated with a type I interferon signature. Nat Genet 44, 1243–1248 (2012).

14. Shallev, L. et al. Decreased A-to-I RNA editing as a source of keratinocytes’ dsRNA in psoriasis. RNA 24, 828–840 (2018).

15. Tossberg, J. T., Heinrich, R. M., Farley, V. M., Crooke, P. S., III & Aune, T. M. Adenosine-to-Inosine RNA Editing of Alu Double-Stranded (ds)RNAs Is Markedly Decreased in Multiple Sclerosis and Unedited Alu dsRNAs Are Potent Activators of Proinflammatory Transcriptional Responses. J Immunol 205, 2606–2617 (2020).

16. Roth, S. H. et al. Increased RNA Editing May Provide a Source for Autoantigens in Systemic Lupus Erythematosus. Cell Reports 23, 50–57 (2018).

17. Vlachogiannis, N. I. et al. Increased adenosine-to-inosine RNA editing in rheumatoid arthritis. Journal of Autoimmunity 106, 102329 (2020).

18. Song, B., Shiromoto, Y., Minakuchi, M. & Nishikura, K. The role of RNA editing enzyme ADAR1 in human disease. WIREs RNA 13, e1665 (2022).

19. Li, Q. et al. RNA editing underlies genetic risk of common inflammatory diseases. Nature 608, 569–577 (2022).

20. Karmon, M. et al. Altered RNA Editing in Atopic Dermatitis Highlights the Role of Double-Stranded RNA for Immune Surveillance. Journal of Investigative Dermatology 143, 933–943.e8 (2023).

21. Knebel, U. E. et al. Disrupted RNA editing in beta cells mimics early-stage type 1 diabetes. Cell Metabolism 36, 48–61.e6 (2024).

22. Tamizkar, K. H. & Jantsch, M. F. RNA editing in disease: mechanisms and therapeutic potential. RNA 31, 359–368 (2025).

23. Wang, Q. D. et al. Stress-induced apoptosis associated with null mutation of ADAR1 RNA editing deaminase gene. Journal of Biological Chemistry 279, 4952–4961 (2004).

24. Hartner, J. C. et al. Liver disintegration in the mouse embryo caused by deficiency in the RNA-editing enzyme ADAR1. Journal of Biological Chemistry 279, 4894–4902 (2004).

25. Li, J. B. & Walkley, C. R. Leveraging genetics to understand ADAR1-mediated RNA editing in health and disease. Nat Rev Genet 26, 532–546 (2025).

26. Athanasiadis, A., Rich, A. & Maas, S. Widespread A-to-I RNA Editing of Alu-Containing mRNAs in the Human Transcriptome. PLOS Biology 2, e391 (2004).

27. Blow, M., Futreal, P. A., Wooster, R. & Stratton, M. R. A survey of RNA editing in human brain. Genome Res 14, 2379–2387 (2004).

28. Kim, D. D. Y. et al. Widespread RNA Editing of Embedded Alu Elements in the Human Transcriptome. Genome Res. 14, 1719–1725 (2004).

29. Levanon, E. Y. et al. Systematic identification of abundant A-to-I editing sites in the human transcriptome. Nature Biotechnology 22, 1001–1005 (2004).

30. Kleinberger, Y. & Eisenberg, E. Large-scale analysis of structural, sequence and thermodynamic characteristics of A-to-I RNA editing sites in human Alu repeats. BMC genomics 11, 453 (2010).

31. Bazak, L. et al. A-to-I RNA editing occurs at over a hundred million genomic sites, located in a majority of human genes. Genome Research 24, 365–376 (2014).

32. Porath, H. T., Carmi, S. & Levanon, E. Y. A genome-wide map of hyper-edited RNA reveals numerous new sites. Nature Communications 5, 4726 (2014).

33. Lander, E. S. et al. Initial sequencing and analysis of the human genome. Nature 409, 860–921 (2001).

34. Batzer, M. A. & Deininger, P. L. Alu repeats and human genomic diversity. Nat Rev Genet 3, 370–379 (2002).

35. Ahmad, S. et al. Breaching Self-Tolerance to Alu Duplex RNA Underlies MDA5-Mediated Inflammation. Cell 172, 797–810.e13 (2018).

36. Sela, N. et al. Comparative analysis of transposed element insertion within human and mouse genomes reveals Alu’s unique role in shaping the human transcriptome. Genome Biology 8, R127 (2007).

37. Piechotta, M., Wyler, E., Ohler, U., Landthaler, M. & Dieterich, C. JACUSA: site-specific identification of RNA editing events from replicate sequencing data. BMC Bioinformatics 18, 7 (2017).

38. Eisenberg, E. & Levanon, E. Y. A-to-I RNA editing - Immune protector and transcriptome diversifier. Nature Reviews Genetics 19, 473–490 (2018).

39. Roth, S. H., Levanon, E. Y. & Eisenberg, E. Genome-wide quantification of ADAR adenosine-to-inosine RNA editing activity. Nature Methods 16, 1131–1138 (2019).

40. Flati, T. et al. HPC-REDItools: a novel HPC-aware tool for improved large scale RNA-editing analysis. BMC Bioinformatics 21, 353 (2020).

41. Lo Giudice, C., et al. Quantifying RNA Editing in Deep Transcriptome Datasets. Front. Genet. 11, (2020).

42. Kluesner, M. G. et al. MultiEditR: The first tool for the detection and quantification of RNA editing from Sanger sequencing demonstrates comparable fidelity to RNA-seq. Molecular Therapy Nucleic Acids 25, 515–523 (2021).

43. Rosenwasser, Z., Levanon, E., Levitt, M. & Oren, G. Detection of RNA Editing Sites by GPT Fine-tuning. in (2024).

44. Torkler, P. et al. LoDEI: a robust and sensitive tool to detect transcriptome-wide differential A-to-I editing in RNA-seq data. Nat Commun 15, 9121 (2024).

45. Eisenberg, E. Chapter Eleven - Bioinformatic approaches for accurate assessment of A-to-I editing in complete transcriptomes. in Methods in Enzymology (ed. Beal, P.) vol. 710 241–265 (Academic Press, 2025).

46. Han, L. et al. The Genomic Landscape and Clinical Relevance of A-to-I RNA Editing in Human Cancers. Cancer Cell 28, 515–528 (2015).

47. Tan, M. Dynamic landscape and regulation of RNA editing in mammals. Nature 550, 249–254 (2017).

48. Breen, M. S. et al. Global landscape and genetic regulation of RNA editing in cortical samples from individuals with schizophrenia. Nat Neurosci 22, 1402–1412 (2019).

49. Ishizuka, J. J. et al. Loss of ADAR1 in tumours overcomes resistance to immune checkpoint blockade. Nature 565, 43–48 (2019).

50. Schaffer, A. A. et al. The cell line A-to-I RNA editing catalogue. Nucleic acids research 48, 5849–5858 (2020).

51. Giorgio, S. D., Martignano, F., Torcia, M. G., Mattiuz, G. & Conticello, S. G. Evidence for host-dependent RNA editing in the transcriptome of SARS-CoV-2. Science Advances (2020) doi:10.1126/sciadv.abb5813.

52. Buchumenski, I. et al. Systematic identification of A-to-I RNA editing in zebrafish development and adult organs. Nucleic Acids Research 49, 4325–4337 (2021).

53. Szymczak, F. et al. ADAR1-dependent editing regulates human β cell transcriptome diversity during inflammation. Front. Endocrinol. 13, (2022).

54. Picardi, E., Mansi, L. & Pesole, G. Detection of A-to-I RNA Editing in SARS-COV-2. Genes 13, 41 (2022).

55. Cuddleston, W. H. et al. Cellular and genetic drivers of RNA editing variation in the human brain. Nat Commun 13, 2997 (2022).

56. Amweg, A. et al. The A to I editing landscape in melanoma and its relation to clinical outcome. RNA Biology (2022).

57. Merdler-Rabinowicz, R., et al. Elevated A-to-I RNA editing in COVID-19 infected individuals. NAR Genomics and Bioinformatics 5, (2023).

58. Avram-Shperling, A., et al. Identification of exceptionally potent adenosine deaminases RNA editors from high body temperature organisms. PLOS Genetics 19, e1010661 (2023).

59. Mann, T. D., Kopel, E., Eisenberg, E. & Levanon, E. Y. Increased A-to-I RNA editing in atherosclerosis and cardiomyopathies. PLOS Computational Biology 19, e1010923 (2023).

60. Rodriguez de los Santos, M., et al. Divergent landscapes of A-to-I editing in postmortem and living human brain. Nat Commun 15, 5366 (2024).

61. Wang, Y. et al. Global characterization of RNA editing in genetic regulation of multiple ovarian cancer subtypes. Molecular Therapy Nucleic Acids 35, (2024).

62. Heraud-Farlow, J. E. et al. GGNBP2 regulates MDA5 sensing triggered by self double-stranded RNA following loss of ADAR1 editing. Science Immunology 9, eadk0412 (2024).

63. Peleg, S. et al. RNA editing deficiency models differential immunogenicity of pancreatic α- and β-cells. Molecular Metabolism 98, 102183 (2025).

64. D’Addabbo, P. et al. REDIportal: toward an integrated view of the A-to-I editing. Nucleic Acids Res 53, D233–D242 (2025).

65. Dong, S. et al. YRA1 Autoregulation Requires Nuclear Export and Cytoplasmic Edc3p-Mediated Degradation of Its Pre-mRNA. Molecular Cell 25, 559–573 (2007).

66. Barak, M. et al. Purifying selection of long dsRNA is the first line of defense against false activation of innate immunity. Genome Biology 21, 26 (2020).

67. Bazak, L., Levanon, E. Y. & Eisenberg, E. Genome-wide analysis of Alu editability. Nucleic Acids Res 42, 6876–6884 (2014).

68. Levanon, E. Y., Cohen-Fultheim, R. & Eisenberg, E. In search of critical dsRNA targets of ADAR1. Trends in Genetics 40, 250–259 (2024).

69. Patterson, J. B. & Samuel, C. E. Expression and Regulation by Interferon of a Double-Stranded-RNA-Specific Adenosine Deaminase from Human Cells: Evidence for Two Forms of the Deaminase. Molecular and Cellular Biology 15, 5376–5388 (1995).

70. van Gemert, F. et al. ADARp150 counteracts whole genome duplication. Nucleic Acids Res 52, 10370–10384 (2024).

71. Colli, M. L. et al. An integrated multi-omics approach identifies the landscape of interferon-α-mediated responses of human pancreatic beta cells. Nat Commun 11, 2584 (2020).

72. Ramos-Rodríguez, M. et al. The impact of proinflammatory cytokines on the β-cell regulatory landscape provides insights into the genetics of type 1 diabetes. Nat Genet 51, 1588–1595 (2019).

73. Gonzalez-Duque, S. et al. Conventional and Neo-antigenic Peptides Presented by β Cells Are Targeted by Circulating Naïve CD8+ T Cells in Type 1 Diabetic and Healthy Donors. Cell Metabolism 28, 946–960.e6 (2018).

74. Kim, J. I. et al. RNA editing at a limited number of sites is sufficient to prevent MDA5 activation in the mouse brain. PLOS Genetics 17, e1009516 (2021).

75. Kleinova, R. et al. The ADAR1 editome reveals drivers of editing-specificity for ADAR1-isoforms. Nucleic Acids Res 51, 4191–4207 (2023).

76. Zaghlool, A. et al. Characterization of the nuclear and cytosolic transcriptomes in human brain tissue reveals new insights into the subcellular distribution of RNA transcripts. Sci Rep 11, 4076 (2021).

77. Lonsdale, J. et al. The Genotype-Tissue Expression (GTEx) project. Nature Genetics 45, 580–585 (2013).

78. Thompson, E. G. et al. Host blood RNA signatures predict the outcome of tuberculosis treatment. Tuberculosis 107, 48–58 (2017).

79. Chen, C.-X. et al. A third member of the RNA-specific adenosine deaminase gene family, ADAR3, contains both single- and double-stranded RNA binding domains. RNA 6, 755–767 (2000).

80. Leinonen, R., Sugawara, H., Shumway, M., & on behalf of the International Nucleotide Sequence Database Collaboration. The Sequence Read Archive. Nucleic Acids Res 39, D19–D21 (2011).

81. Barrett, T. et al. NCBI GEO: archive for functional genomics data sets—update. Nucleic Acids Research 41, D991–D995 (2013).

82. Mailman, M. D. et al. The NCBI dbGaP database of genotypes and phenotypes. Nat Genet 39, 1181–1186 (2007).

83. Weinstein, J. N. et al. The Cancer Genome Atlas Pan-Cancer analysis project. Nat Genet 45, 1113–1120 (2013).

84. Heath, A. P. et al. The NCI Genomic Data Commons. Nat Genet 53, 257–262 (2021).

85. Kent, W. J. et al. The Human Genome Browser at UCSC. Genome Research 12, 996–1006 (2002).

86. Quinlan, A. R. & Hall, I. M. BEDTools: A flexible suite of utilities for comparing genomic features. Bioinformatics 26, 841–842 (2010).

87. Patro, R., Duggal, G., Love, M. I., Irizarry, R. A. & Kingsford, C. Salmon provides fast and bias-aware quantification of transcript expression. Nat Methods 14, 417–419 (2017).

88. Dobin, A. et al. STAR: ultrafast universal RNA-seq aligner. Bioinformatics 29, 15–21 (2013).

89. Chen, S., Zhou, Y., Chen, Y. & Gu, J. fastp: an ultra-fast all-in-one FASTQ preprocessor. Bioinformatics 34, i884–i890 (2018).

90. Sherry, S. T. et al. dbSNP: the NCBI database of genetic variation. Nucleic Acids Res 29, 308–311 (2001).

91. Gabay, O. et al. Landscape of adenosine-to-inosine RNA recoding across human tissues. Nature Communications 2022 13:1 13, 1–17 (2022).

92. Pinto, Y., Cohen, H. Y. & Levanon, E. Y. Mammalian conserved ADAR targets comprise only a small fragment of the human editosome. Genome biology 15, R5 (2014).

93. Li, H. et al. The Sequence Alignment/Map format and SAMtools. Bioinformatics 25, 2078–2079 (2009).

94. Dale, R. K., Pedersen, B. S. & Quinlan, A. R. Pybedtools: a flexible Python library for manipulating genomic datasets and annotations. Bioinformatics 27, 3423–3424 (2011).

95. Kaplan, E. L. & Meier, P. Nonparametric Estimation from Incomplete Observations. Journal of the American Statistical Association 53, 457–481 (1958).

96. Di Tommaso, P. et al. Nextflow enables reproducible computational workflows. Nat Biotechnol 35, 316–319 (2017).

97. Hulsen, T. DeepVenn -- a web application for the creation of area-proportional Venn diagrams using the deep learning framework Tensorflow.js. Preprint at 10.48550/arXiv.2210.04597 (2022).

